# What are the best practices for curating eDNA custom barcode reference libraries? A case study using Australian subterranean fauna

**DOI:** 10.1101/2024.09.18.611555

**Authors:** Michelle T. Guzik, Danielle N. Stringer, Jake Thornhill, Peterson J. Coates, Mieke van der Heyde, Mia J. Hillyer, Nicole E White, Mattia Saccò, Perry Beasley-Hall, William F. Humphreys, Mark S. Harvey, Joel A. Huey, Nerida G. Wilson, Jason Alexander, Garth Humphreys, Rachael A. King, Steve J. B. Cooper, Adrian Pinder, Giulia Perina, Andrew M. Hosie, Lisa Kirkendale, Paul Nevill, Andy D. Austin

## Abstract

Identification of species for environmental assessment and monitoring is essential for understanding anthropogenic impacts on biodiversity, but for subterranean fauna this task is frequently difficult and time consuming. The implementation of environmental DNA (eDNA) metabarcoding for biodiversity discovery and assessment offers considerable promise for improving the rate, accuracy and efficiency of species detection in ecosystems both above and below the ground. Importantly, for a better understanding of the biodiversity and ecology of organisms detected using eDNA, a custom library of known reference sequences with associated correct taxonomic metadata—i.e., a barcode reference library (BRL)—is required. Yet, minimal guidance is currently available on how an effective (i.e. shareable, multi-sequence, that permits metadata and has a unified nomenclature) and accurate (i.e. verified) custom BRL can be achieved. Here, we present a detailed roadmap for curation of a BRL for subterranean fauna. To do this, we (1) curated a custom sequence database of subterranean fauna at an environmentally sensitive location, Bungaroo Creek in the Pilbara region of Western Australia, for four gene loci useful for eDNA metabarcoding (*COI*, *18S* rRNA, *12S* rRNA and *16S* rRNA); (2) addressed major gaps in taxonomy and disparate nomenclature of subterranean fauna by estimating 17–29 putative new species with standard delimitation methods, including 34 Barcode Index Numbers (BINs) in BOLD, and (3) summarised a best practice workflow for curation of a custom BRL that has broad applicability and can be applied to any taxa.

**Scientific Significance Statement:** In threatened ecosystems, environmental DNA (eDNA) metabarcoding for biodiversity discovery and assessment offers considerable promise for improvement in the rate, efficiency and accuracy of species detection. For a better understanding of the biodiversity and ecology of organisms detected using eDNA, a custom library of known reference sequences with associated correct taxonomic metadata is required. Minimal guidance is currently available on how an effective (i.e. shareable, multi-sequence, permits metadata and provides a unified nomenclature) custom barcode reference library (BRL) can be achieved for subterranean fauna. Here, we present a road map for sound and reliable curation of a BRL using subterranean fauna from Australia as a case study.

## Introduction

Groundwater is a high-value natural resource that is vulnerable to human-induced impacts and climate change (Kuang *et al*. 2024). In Australia, the economic value of groundwater resources is estimated to be $33.8 billion (Deloitte Access Economics 2013). Importantly, groundwater also supports groundwater-dependent ecosystems (GDEs), such as subterranean habitats, where a unique and ancient fauna provide essential ecosystem services that maintain underground water condition and quality (Boulton *et al*. 2008). Globally, considerable tension exists with respect to the use of groundwater for economic growth and environmental sustainability (Mammola *et al*. 2019; Saccò *et al*. 2022a; Mammola *et al*. 2024; Saccò *et al*. 2024). In certain countries, groundwater fauna are surveyed during routine environmental impact assessments (EIAs) using biomonitoring methods (Environmental Protection Authority 2021), particularly in areas of potential resource development (Gibson *et al*. 2019). For all stakeholders reliant on such biomonitoring, four important issues frequently impact decision-making capabilities (Gibson *et al*. 2019; Mammola *et al*. 2021; Saccò *et al*. 2022b): 1) inconsistent metadata quality (Costello *et al*. 2013; Saccò *et al*. 2022b); 2) minimal data sharing (e.g., Meyer *et al*. (2015)); 3) uncertainty around sampling effort required to provide a robust understanding of the values and predicted impacts and 4) high numbers of undescribed species (Costello 2015; Ficetola *et al*. 2015) with insufficient associated taxonomic knowledge (Mammola *et al*. 2019; Mammola *et al*. 2021). Overcoming these impediments is complicated by reduced taxonomic capacity for many groups and difficulty in distinguishing species on morphological characteristics alone due to anatomical convergence and their often-cryptic nature (Abrams *et al*. 2012; King *et al*. 2022).

Environmental DNA (eDNA) metabarcoding has been identified as an approach with significant potential to alleviate the difficulties associated with traditional sampling (van der Heyde et al. 2023). Indeed, it has been demonstrated as effective for detection of subterranean fauna (West *et al*. 2020; White *et al*. 2020; van der Heyde *et al*. 2023a; van der Heyde *et al*. 2023b). To identify subterranean fauna using eDNA, sequences belonging to anonymous operational taxonomic units (OTUs) must be assigned taxonomic information from a reference database (Taberlet *et al*. 2012). However, this assignment is often patchy due to a lack of data in public DNA databases (e.g., BOLD and GenBank) for both genes and taxonomically obscure species as well as inconsistencies in taxonomic metadata availability (Costello 2015; Kvist 2013; Dormontt *et al*. 2018; DeWaard *et al*. 2019; Ficetola *et al*. 2019). For these reasons, much of the eDNA data being generated by metabarcoding studies are likely underutilised (Korbel *et al*. 2017). While taxonomy-independent approaches to eDNA bioinformatics are being developed (Brinkmann *et al*. 2023; Wilkinson *et al*. 2024), many legislative frameworks for subterranean fauna require species-level identifications for bioassessment, although more holistic approaches are being advocated (Koch *et al*. 2024).

Therefore, custom reference databases for querying OTUs are critical for the successful implementation of eDNA metabarcoding methods (Saccò *et al*. 2022b) and resulting biodiversity assessments. This is especially the case given the extent of short-range endemism as well as the numerous threatening process that impact subterranean communities (Harvey *et al*. 2011). Guidance on how these databases should be curated is sorely needed.

Selection criteria for the curation of specimens and DNA sequences when submitting to public databases are established for certain taxonomic groups of subterranean fauna (pseudoscorpions (Harvey *et al*. 2023; Hlebec *et al*. 2023); isopods and myriapods (Rendoš *et al*. 2023); asellid isopods (Saclier *et al*. 2024)). Such criteria include: morphologically verified specimens; homologous DNA sequences corresponding to common metabarcoding assay regions; estimating levels of genetic variation within and between species; and detailed metadata, linking specimens and current taxonomic information per sequence.

Problematically, for understudied and biodiverse groups such as invertebrate fauna, sequence records frequently lack associated taxonomic information as it often does not exist. Instead, a unified system of nomenclature that does not solely rely on formally described species is required. Various methods for delimiting species are already widely used (e.g., Automatic Barcode Gap Discovery (ABGD; Puillandre *et al*. (2012)), Assemble Species by Automatic Partitioning (ASAP; Puillandre *et al*. (2021)) and phylogenetic methods such as Bayesian Poisson Tree Processes (bPTP; Zhang *et al*. (2018)). Such methods require specialist knowledge to implement, and are frequently not linked to an existing database platform that is user-friendly, and where the sequences are available in the same place. Whilst some authors have suggested that these approaches be used as an alternative to formal taxonomic approaches (Meierotto *et al*. 2019; Sharkey *et al*. 2021), most researchers recognise that a major problem exists in providing up-to-date and shareable nomenclature in the absence of taxonomic information.

To combat the taxonomic impediment (Engel *et al*. 2021) facing subterranean fauna (Mammola *et al*. 2021), delimited lineages may act as a proxy for species until such time as these species are formally described. The Barcode of Life Database (BOLD) applies a system of Barcode Index Numbers (BINs), which are an index of unique identifiers based on the sequences of varying length and specimens that exist in BOLD. The BIN pipeline analyses new sequence data for the cytochrome *c* oxidase subunit I (*COI*) barcoding gene as they are uploaded to the database platform. The power of the BIN system is that it provides a unified nomenclature for sequences entirely independent of the sequence submitter and their taxonomic capacity, and that information is retained as part of the database. Here, we present a roadmap for curation of a custom barcode reference library (BRL) for subterranean fauna using the BOLD system and build on the new and rapidly expanding area of eDNA metabarcoding research. Our roadmap focuses on an exemplar groundwater-dependent ecosystem at Bungaroo Creek, located in the resource-rich Pilbara region of Western Australia (WA). In future, this data infrastructure can be expanded across Australia and globally in subsequent research since it meets critical principles of a findable, accessible, interoperable and reusable (FAIR) system of data management outlined by Wilkinson *et al*. (2016).

Subterranean biodiversity is comprehensively understudied (Guzik *et al*. 2011), but depending on the region, subterranean fauna can be repeatedly monitored or remain completely unknown to science (Saccò *et al*. 2022a). Given the articulated need for a subterranean fauna BRL by multiple stakeholders (i.e. industry, regulators and academic researchers (Gibson *et al*. 2019)), our overarching aim was to develop a protocol that is easily accessible and usable for effective and efficient characterisation of subterranean biodiversity.

Our aims were to:

1. Curate a *de novo* custom BRL of sequences for an exemplar subterranean faunal community of high biodiversity value.
2. Investigate a unified nomenclature that is stable and repeatable, even in the absence of formal description of species, by using species delimitation methods, such as BINs, to identify and name species proxies.
3. Develop and implement a best-practice workflow for a future dynamic subterranean fauna BRL. We outline a protocol for generating and uploading DNA sequence data and associated metadata for the exemplar location using preferred criteria (i.e. 1–2 listed above).

## Methods

### Bungaroo Creek, WA: a high biodiversity value subterranean community

The Pilbara region of WA is globally recognised for its significant subterranean fauna (Halse *et al*. 2014; Deharveng *et al*. 2024), with many distinct and isolated communities. The Robe River supports several subterranean communities of high ecological value, including that below the major tributary known as Bungaroo Creek (Clark *et al*. 2021). Featuring channel iron deposits and alluvial aquifers, this section of the Robe River provides an ideal habitat for subterranean fauna. Indeed, evolutionarily distinct taxon groups such as the blind cave eel (*Ophisternon candidum*), hydrobiid snails, copepods, ostracods, bathynellaceans and unique amphipod lineages make this environmentally complex location a significant subterranean community (Clark *et al*. 2021) which is recognised as a Priority Ecological Community (Priority 1) in Western Australia (Department of Biodiversity, Conservation and Attractions 2023). Due to the use of Bungaroo Creek groundwater to supply water to the West Pilbara Water Supply Scheme, the subterranean community is frequently sampled, surveyed, and monitored for assessment of environmental impact. Here, we used the subterranean community present at Bungaroo Creek as an exemplar location for the curation of a custom reference library of DNA sequences for future eDNA metabarcoding.

### Specimen collection and curation

To curate a *de novo* custom reference library of subterranean fauna from Bungaroo Creek (herein called Bungaroo BRL), whole specimens from 21 mining exploration bores were obtained for Sanger sequencing (Figure 1). Initially, samples were obtained from archival collections of subterranean fauna from surveys undertaken during 2014 and 2017 and deposited at the Western Australian Museum (WAM). Collection methods included net hauling (Saccò *et al*. 2022b) from aquifers using pre-existing bore holes. Morphological identification was undertaken to the lowest possible taxonomic level, with some specimens identified to morphospecies but most only to order level or higher.

**Figure 1:**
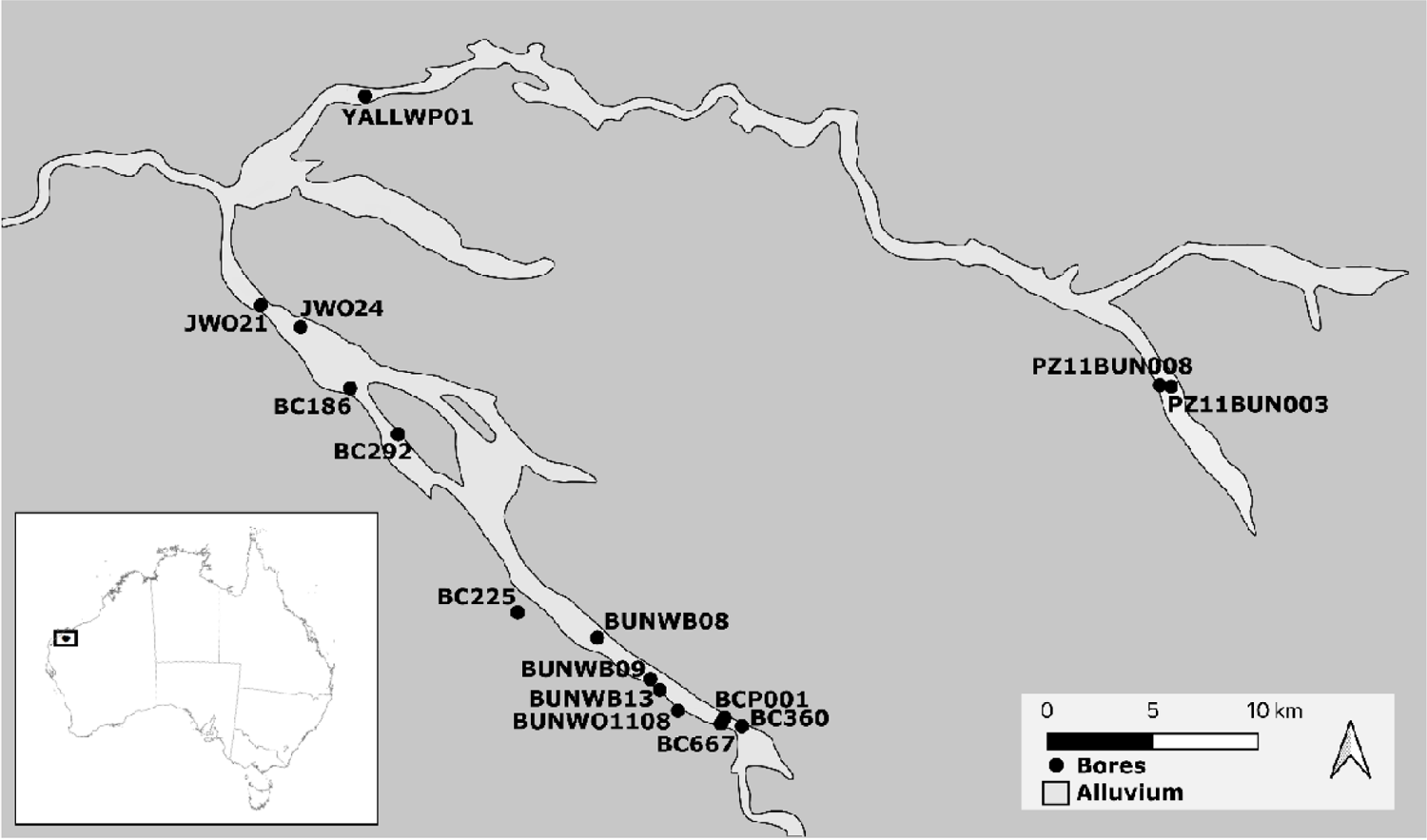
Map showing the locations of sampled boreholes at the Bungaroo Creek site, with bore locations and names. Map created using ArcMap v 10.3.1 (ESRI 2017), data layers provided by Rio Tinto.

In addition to the archived material from the WAM collection, invertebrates were also sampled at the Bungaroo Creek borefield site during June 2017 (as above). Bores were selected using previous survey data with the aim of capturing the greatest possible genetic diversity within the constraints of time and logistical feasibility. The locations of bores sampled are provided in Figure 1, Supplementary Material 1. Invertebrate specimens were taken from each bore using stygofauna haul nets. Five consecutive haul net samples were collected, each involving lowering of a net to the bottom of the bore, with nets being sterilised using bleach between sites to prevent contamination. Bulk invertebrate samples were stored in a portable refrigerator until the end of sampling each day and were then transferred to a 4°C refrigerator. The samples for each bore were pooled, and specimens were sorted *in situ* into major taxonomic groups using a dissecting microscope. Specimens of the same order were each stored in glass vials, labelled with a single unique identifier, and stored at –20°C in 100% ethanol until DNA extraction. Representative individuals were catalogued at the WAM for reference (see Supplementary Material 1 for catalogue information metadata).

### Barcode gene regions

Barcode gene regions are specific segments of DNA sequences that are utilised for molecular taxonomic identification, particularly in the context of delineating species. This was first proposed in 2003 (BOLD; Hebert *et al*. (2003)) where standardised DNA sequences were employed to identify different species. The barcode gene regions selected for this purpose are typically short (∼600-700bp) containing highly conserved regions to allow for primer binding, which then amplify across a variable region to allow for the differentiation of species at the molecular level. The most commonly used barcode region for animals is *COI*, but other genes are also used (https://v3.boldsystems.org/index.php/resources/boldfaq). In order to curate a useable custom library for Bungaroo Creek, we sequenced three mitochondrial genes (*COI*, *12S* and *16S*) and one nuclear gene (*18S*). Therefore, we ensured our BRL would; 1) allow for a strong match to existing metabarcoding assay primers and provide a reference for the development of new metabarcoding primers (if required); 2) ensure accuracy and breadth of species identification and documentation of biodiversity (Epp *et al*. 2012; Hebert *et al*. 2013; Deagle *et al*. 2014); and 3) allow the consolidation of sequence data from other database libraries, both old and new.

### DNA extraction, amplification, and sequencing

DNA was extracted from 468 specimens in December 2017 using a Qiagen DNeasy Blood and Tissue Kit. Established universal invertebrate primers (below) were used for PCR and sequencing of specimens, but optimisation of PCR conditions were minimally adjusted between specimens (Supplementary Material 2). Products were visualised using the eGel Electrophoresis system (Life Technologies, Scoresby, Victoria, Australia) on 2% agarose gels with ethidium bromide. Bidirectional long-read Sanger sequencing was carried out at the Australian Genome Research Facility (Adelaide).

### DNA sequence database submissions

For specimens from the Bungaroo BRL, haplotype sequences new to this study were submitted to GenBank (accession numbers: OR524806-OR524929) and BOLD (public BOLD Project: BUBRL https://boldsystems.org/index.php/MAS_Management_DataConsole?codes=BUBRL) (Supplementary Material 3) including near-complete taxonomy and provenance of the data. Our rationale for curating and submitting data to both public platforms was that, while GenBank is the most ubiquitously used to query metabarcoding data due to the BLAST function, BOLD has many of the GUI features for curating and storing metadata needed for the current study and any future work. Specifically, BOLD is a long-term funded, user-friendly web interface with taxonomic, geographic, and photographic metadata available for each sequence (Ratnasingham and Hebert 2007). In addition, BOLD hosts an *in silico* delimitation algorithm that assigns identifiers (BINs) to *COI* lineages based on the genetic distance between all sequences in its database. The BINs provide a centralised system for identifying lineages that are potentially equivalent to species (Hajibabaei *et al*. 2007). While BINs frequently do not exist for underrepresented taxonomic groups and regions (Lahaye *et al*. 2008), we consider that this study fills in some of the gaps in nomenclature and taxonomy that substantially impact data sharing and transferability between projects. GenBank lacks most of these features and, because it does not require the association of sequence data with specimen vouchers, the taxonomy associated with a sequence can never be verified.

### Nucleotide analysis and species delimitation

Geneious v10.0.9 (https://www.geneious.com) was used to edit and align *COI*, *12S, 16S* and *18S* sequence data for all individuals. Forward and reverse sequence fragments were aligned, edited and trimmed. *COI* sequencers were translated into amino acids to ensure no stop codons and a single consensus sequence was created. Consensus sequences of all individuals for each gene were aligned using MUSCLE (Edgar 2004), trimmed to equal lengths, and FaBox (Villesen 2007) was used to collapse the final sequence alignments into haplotypes using its *DNA to haplotype collapser and converter* functionality. Automatic Barcode Gap Discovery (ABGD) (Puillandre *et al*. 2012), a computationally efficient distance-based method for species delimitation, was implemented to identify putative species (lineages equivalents) (https://bioinfo.mnhn.fr/abi/public/abgd/abgdweb.html). This method seeks to quantify the location of the barcode gap that separates intra-from inter-specific distances. It performs well when compared to tree-based coalescent methods (Puillandre *et al*. 2012; Kekkonen and Hebert 2014; Kapli *et al*. 2017) and other threshold techniques (Ratnasingham and Hebert 2013).

Further to ABGD, we implemented the BOLD BIN method of lineage delimitation. BINs are an index of unique identifiers based on a database of specimens belonging to each BIN with their associated metadata as implemented by BOLD systems. The pipeline analyses new sequence data for the *COI* barcoding region as they are uploaded to BOLD by applying a two-stage algorithm (Refined Single Linkage) that couples single linkage and Markov clustering to assign sequences to a sequence cluster and is subsequently assigned a unique identifier termed a BIN (Ratnasingham and Hebert 2013). The Refined Single Linkage algorithm matches the taxonomic performance of competing approaches, but couples this with protocols that are simple enough to allow the automated assignment of all new barcode records to a BIN. Sequences that establish a new BIN add an entry to the BIN index, while sequences assigned to an existing BIN contribute their metadata to it.

## Results

### Summary of haplotypes and genetic distances

A total of 468 individuals were available for PCR and sequencing, but success rates for the total suite of specimens were low (*COI*: *n* = 103 (22%), *18S*: *n* = 137 (29%), *12S*: *n* = 88 (19%) and *16S*: *n* = 21 (5%)) (see Discussion for explanation). Sequence data were obtained from a total of 195 individuals (42%), including 660 base pairs (bp) for *COI* (*n* = 103 and *h* = 59), 1017 bp for *18S* (*n* = 136 and *h* =58), 425 bp for *12S* (*n* = 88 and *h* = 48), and 511 bp for *16S* (*n* = 21 and *h* = 11). Only seven individuals were successfully sequenced for all four barcoding loci (see Supplementary Material 1). All 195 sequences were submitted to our BOLD project ID BUBRL (Bungaroo BRL) for four barcode genes *COI*-5P (103 sequences), *12S* (88), *16S* (21), and *18S* (136), (GenBank accession numbers *COI*: OR524806 - OR524908; *18S*: OR524930 - OR525066; *12S*: OR525144 - OR525231; *16S*: OR524909-OR524929) and BOLD (public BOLD Project: https://boldsystems.org/index.php/MAS_Management_DataConsole?codes=BUBRL).

### Species delimitation

Haplotype groups equivalent to lineages were delineated using ABGD (Figure 2 and Supplementary Material 4) and compared to BOLD BINs as well as to morphospecies identified by Bennelongia Ltd, an environmental consulting company that undertook routine biomonitoring in the area during a 2017 survey at Bungaroo Creek (Table 1). Results of the ABGD analysis revealed 29 lineages for *COI*, 16 lineages for *12S,* 9 lineages for *16S* and 17 lineages for *18S*. Thirty-four BINs were identified for *COI*. Delineated lineages based on ABGD (for all four genes) and BOLD BINs (based on *COI* only) for major taxon groups included amphipods (ABGD = 4–8 lineages; BINs = 9), bathynellaceans (ABGD = 1–4 lineages; BINs = 4), thermosbenaceans (ABGD = 1–2 lineages; BINs = 3), annelids (ABGD = 1–6 lineages; BINs = 5), ostracods (ABGD = 1–3 lineages; BINs = 3) copepods (ABGD = 4–5 lineages; BINs = 8), and a single lineage of gastropod. It should be noted that Pauropoda are most likely troglophilic/ edaphobitic but were retained here as representatives of the subterranean ecosystem at Bungaroo Creek.

**Figure 2:**
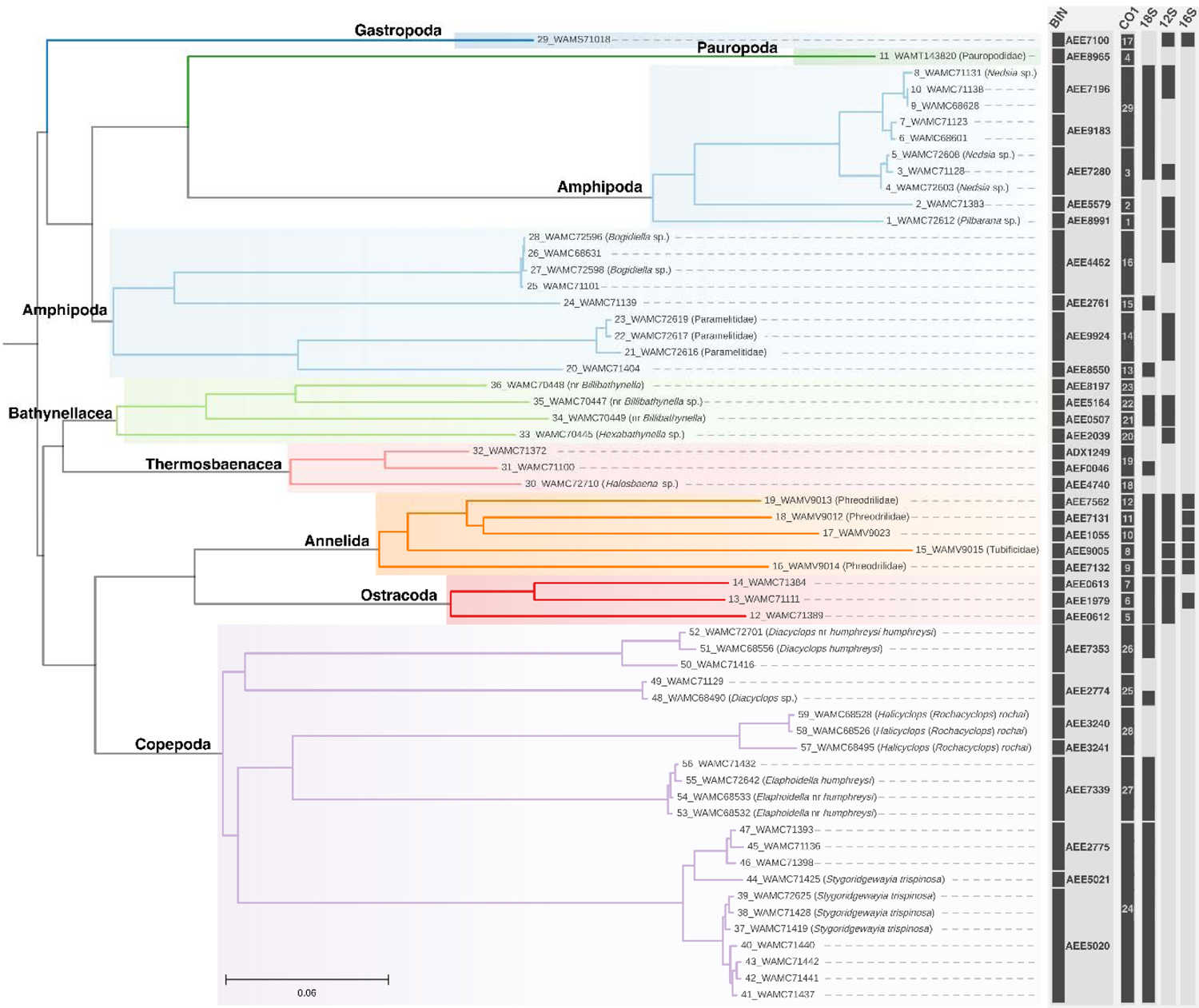
Phylogenetic tree of taxa from Bungaroo creek for the mitochondrial *COI* gene– with species delimitation results represented by taxonomic group in colour: Amphipoda (blue), Copepoda (purple), Bathynellacea (green), Thermosbaenacea (pink), Ostracoda (red), Annelida (orange), Gastropoda (dark blue). Relationships between major taxonomic groups are not representative of actual phylogenetic relationships.

**Table 1:**
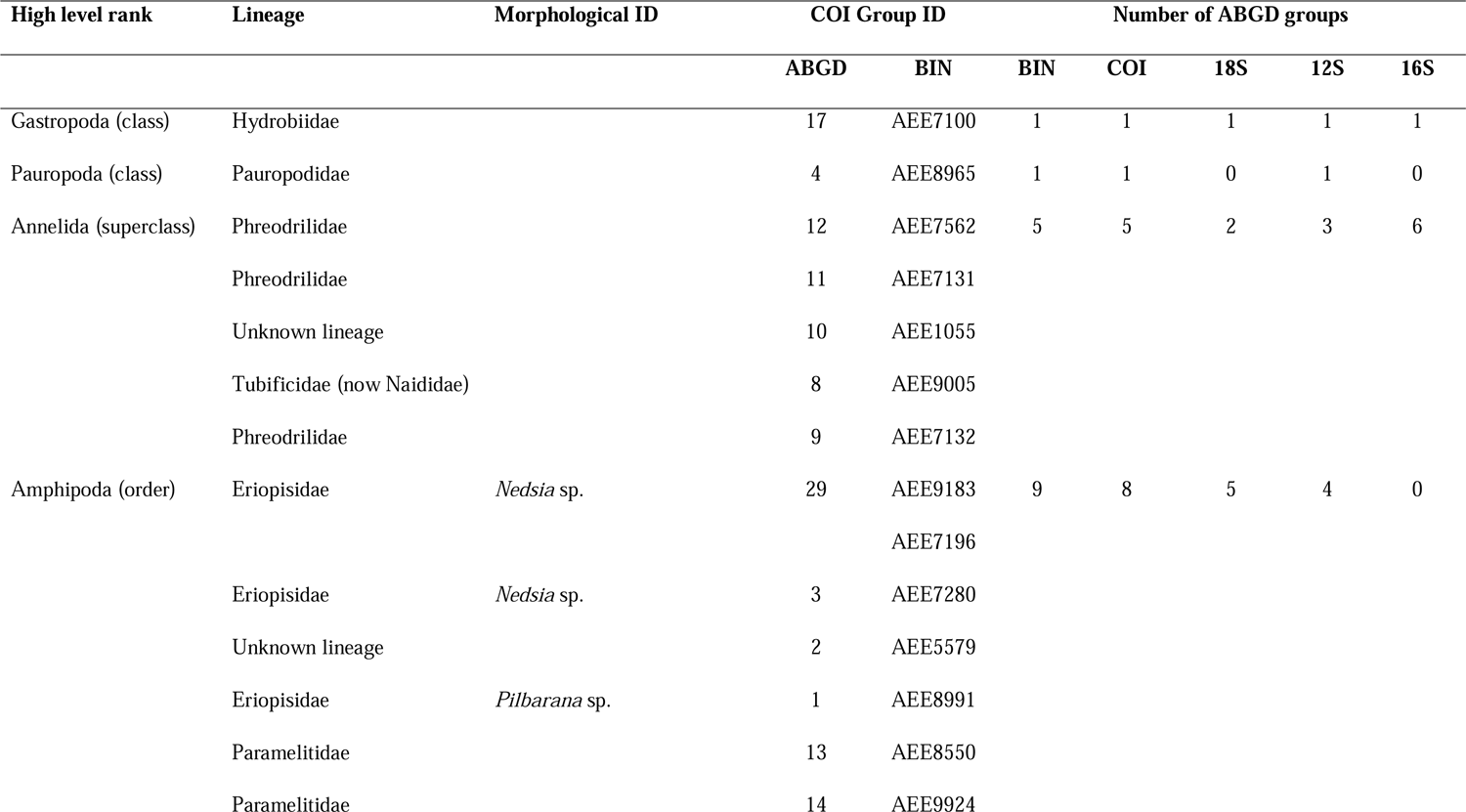
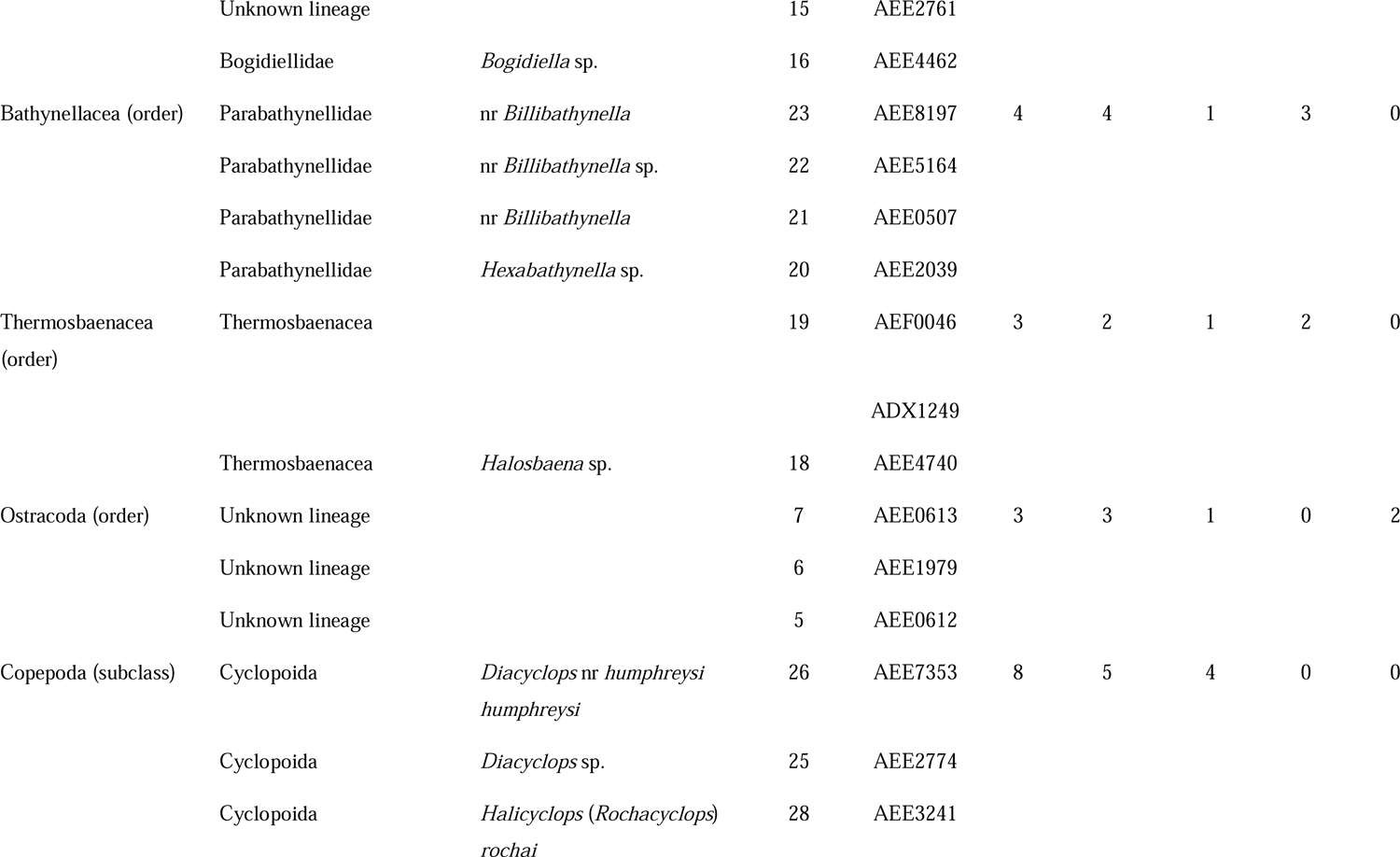
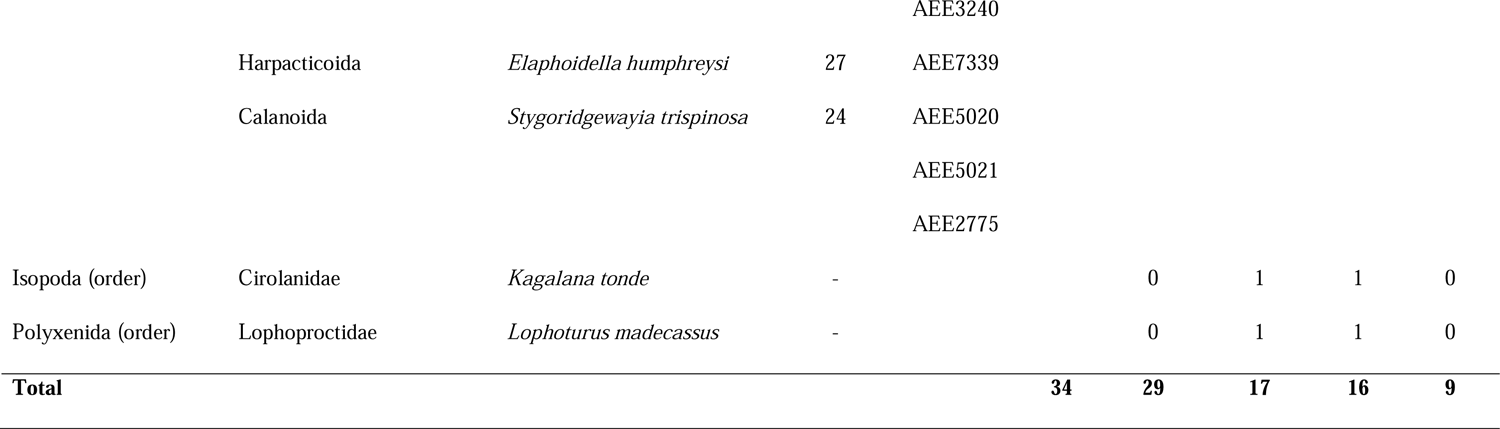
Morphospecies and delimited lineages of subterranean fauna from Bungaroo Creek, Western Australia (for lineages see Figure 2) using BOLD BINs for all four barcoding genes examined in this study.

## Discussion

The use of eDNA metabarcoding to estimate the biodiversity of subterranean faunal communities has the capacity to alleviate a bottleneck that morphological examination of physical specimens, limited taxonomic expertise, low-quality or non-existent taxonomic keys, and a poorly described fauna have on biodiversity assessment (Gibson 2019). The most efficient way to identify species from anonymous eDNA metabarcoding OTUs is to employ a curated reference library of sequences (barcodes), preferably with matching voucher specimens and/or taxonomic information (i.e., a custom BRL). Here, we developed preferred sequence data and metadata criteria for submission to BOLD for subterranean fauna. This best-practice protocol addresses a suite of issues specific to subterranean fauna by applying a unified nomenclature to the exceptional and under-described biodiversity that exists in groundwater-dependent ecosystems, such as the exemplar site chosen for this study. Further, our Bungaroo BRL forms the basis of a publicly available data resource that can be used in eDNA based studies of subterranean fauna into the future.

### Estimating community diversity at Bungaroo Creek

Combined, the four genes sequenced here for Bungaroo Creek subterranean fauna delineated up to 29 genetic lineages as estimated with ABGD based on the *COI* data for 17 taxon groups. The majority of these lineages were further supported by ABGD results for the other three genes. These putative species included mites, amphipods, rotifers, copepods, isopods, worms, snails, and millipedes, indicating a substantially speciose invertebrate community.

The four genes were effective for sequencing the key taxon groups present, but there was some variation in the number of delineated lineages between genes. For instance, *COI* delineated the highest number of lineages (29), followed by *18S* (17), *12S* (16) and *16S* (9). Some of the variation in numbers of delineated species was attributable to sequencing success or lack thereof (e.g. *16S*, see below) (e.g. Asmyhr and Cooper (2012)), as well as a wide range of resolution given the diversity of markers employed, but the delineation methods used here may also reflect strong phylogeographic variation and structuring (Ortiz and Francke 2017).

Compared to other studies, 29 lineages of subterranean fauna represent a significantly more diverse community at Bungaroo than for other Australian aquifers, although this number is consistent with biodiversity assessment surveys based on morphological examinations (e.g. Bennelongia (2013)). For example, we observed between two and three lineages of ancient Thermosbaenacea, whereas only one species is currently described from the Pilbara (Poore and Humphreys 1992; Eberhard *et al*. 2005). Our results are supported by (Page *et al*. 2018) who also detected a suite of genetic lineages indicating a much higher species diversity of this tethyan crustacean than previously thought. In other Australian aquifers, estimates of 8– 16 putative species have been reported for eastern Australian alluvial aquifers (Asmyhr and Cooper 2012), and calcrete aquifers at Sturt Meadows (Bradford *et al*. 2014) and Lake Violet (Watts and Humphreys 2003; Eberhard *et al*. 2005) in the Yilgarn region WA, and the Millstream aquifer in the Pilbara (Eberhard *et al*. 2005). As such, Bungaroo Creek represents one of the highest putative species estimates for an Australian aquifer. Alongside our own study, recent studies by (Perina *et al*. 2018; Perina *et al*. 2019; Matthews *et al*. 2020) reinforce the view that the Pilbara harbours an exceptionally high and little-explored diversity of subterranean faunal species.

### BINs

Barcode Index Numbers (BINs) from genetic databases (i.e., BOLD) have been promoted to offer a standardised and globally unique identifier for each species or species proxy, enhancing comparability and repeatability of studies, allowing researchers to pool data and conduct meta-analyses (Hajibabaei *et al*. 2007; Ratnasingham and Hebert 2007; Lahaye *et al*. 2008). In particular, BINs, based on the genetic distance between sequences independent of prior taxonomic knowledge, have the potential to help identify cryptic or overlooked species to provide an objective means for identifying species boundaries (Hajibabaei *et al*. 2007).

Here, ABGD and BIN estimates of putative species for the Bungaroo BRL were generally consistent, although for the copepods three extra lineages were identified using the latter system. While in some cases BINs have been recognised to have some detrimental aspects in certain cases. For example, over-splitting and potential misidentification of lineages is recognised as problematic (Deagle *et al*. 2014) (i.e. BIN assignment has an 86% success rate (Hartop *et al*. 2022)). Further, for unsampled taxon groups and geographic locations where representation in BOLD is absent, existing BINs do not exist, leading to the creation of many new BINs when sampling increases (Lahaye *et al*. 2008). Finally, BIN assignments can change over time as new sequences and taxon groups are added. However, for large, well-represented BINs, the changes are often small and can be tracked by BOLD (Ratnasingham and Hebert 2013). An important advantage of BINs is their usefulness in providing a unified nomenclature that is publicly available and usable. If required, BINs can be verified using current taxonomy or alternative species delimitation methods (i.e., via morphological and/or molecular methods) and updated on BOLD if needed.

### Recommended workflow for Barcode Reference Library sequence/data additions

Using Bungaroo Creek as an example, we have demonstrated how to create a dynamic subterranean faunal reference library for ongoing and future research and biomonitoring that can be updated and, importantly, follows FAIR Principles of Data Infrastructure (Wilkinson et al. 2016). Below we make recommendations for a suitable workflow for addition of data to our BRL for subterranean fauna (i.e. illustrative summary, Figure 3).

**Figure 3:**
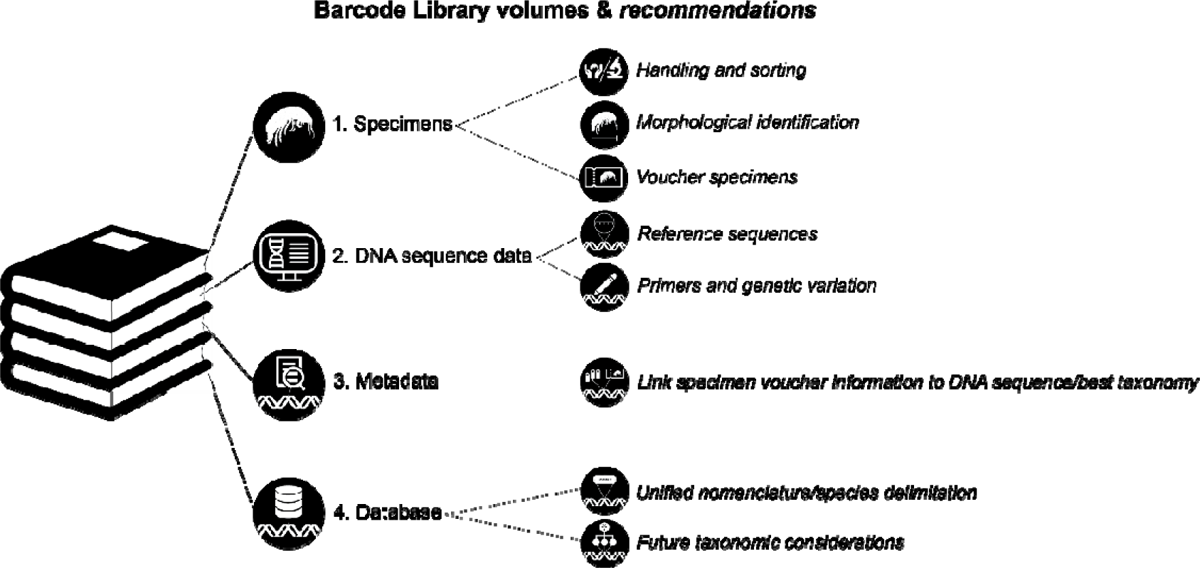
Recommended workflow for construction of a Barcode Reference Library.

### SPECIMENS

#### Handling and sorting: in-field and laboratory

Animal sampling requires compliance with regional collection standards and permits. Tissues should be handled according to those standards. Here, we address handling of invertebrates and their tissue specifically since sampling of subterranean vertebrate specimens pose substantially different requirements (Saccò et al. 2022b).

Development of a BRL first requires long-strand sequencing of DNA (Sanger or next generation sequencing (NGS)) from reference specimens that are also suitable for morphological examination either pre- or post-sequencing. Overall, the quality of specimen preservation for DNA sequencing is critical to the successful development of a reference library. Snap-freezing of specimens is the most optimal for long-term storage of DNA but difficult to achieve in the field, especially at remote locations. Instead, under ideal conditions, specimens should be kept alive and then killed by sorting into cold 100% undenatured ethanol or propylene glycol with no other additives for preservation, and them immediately refrigerated at a minimum of –20° C (Moreau *et al*. 2013). It should be noted that for small specimens, especially microcrustaceans, sorting in a laboratory with good lab practises, ergonomic consideration to prolonged microscopy and quality equipment can yield improved results. Storage of specimens at room temperature is not ideal but may be unavoidable (Nichols *et al*. 2020). For many surveys, it is not feasible to spend extended time in the field, and bulk samples are frequently transported back to the laboratory for sorting. In these situations, it is recommended that the maximum amount of water from bulk samples be removed prior to sorting, with animals sorted live where possible, and the organic material is refrigerated to the lowest temperature possible in 100% ethanol and sorted as soon as possible (Hajibabaei *et al*. 2012). Clean specimen handling is important, and should be undertaken with gloves and instruments cleaned in 100% ethanol to control for cross contamination between specimens. It should be noted that NGS approaches are significantly increasing the potential of using DNA information from degraded specimens in biodiversity analysis (Hajibabaei *et al*. 2012).

#### Morphological identification

For critical taxonomic metadata and establishment of specimen vouchers, morphological analysis of specimens should always be undertaken when adding data to a BRL. A total evidence approach (with morphological and DNA sequence data) permits validation of identified species (whether described or new) and can ultimately be used to establish formal nomenclature for species. While it is important to remember that any species concept is a working hypothesis, we specifically note that morphological characters alone do not always reveal the full extent of species-level genetic diversity in a population or at a location (King *et al*. 2012; King *et al*. 2022). Additionally, it may not always be feasible to identify specimens to species or even genus level due to a lack of taxonomic expertise and/or ongoing funding, the presence of cryptic species, inadequate taxon sampling, the collection of various life-history stages or damaged specimens that make taxonomic identification difficult, and taxonomic uncertainty and/or incompleteness, especially in poorly-explored geographic locations. High-quality images of putative morphotypes taken prior to DNA extraction can be used as a record of morphological features in the event of destructive sampling and can be uploaded to BOLD. While new techniques are being developed to retain the exoskeleton of specimens during DNA extraction and library preparation (e.g. acoustic droplet ejection (Cain-Hom *et al*. 2016)), currently this technology is highly specialised and expensive to implement. However, ongoing development by major consortia, such as the BIOSCAN project (Park *et al*. 2023), has the potential to expand the application of these methods and decrease the cost of widespread implementation. In the interim, intact mature animals should ideally be morphologically identified where possible prior to sequencing to yield morphological and molecular data linked within the same specimen.

#### Voucher specimens

Retention of specimen vouchers for future morphological validation, and description is vital, and further DNA sequencing can aid with verification of genetic lineages. Archived specimens and tissues are valuable as a temporal genetic and morphological record of the status of a species. Specimen storage (Burrell *et al*. 2015) and age of specimens (Miller *et al*. 2013) are important parameters to successful DNA extraction and PCR amplification and can significantly affect the task of building a BRL (Baird *et al*. 2011; Hebert *et al*. 2013; Burrell *et al*. 2015). In order to retain specimens, dissection and extraction of DNA from a part of the body that will minimally interfere with morphological identification is ideal and usually feasible. However, for very small individuals, sequencing of one to five representatives from a group of morphologically identical individuals collected from the same sampling site allows remaining individuals to be treated as a voucher specimen. Recent technical developments have enabled whole genome sequencing of single individuals for tiny species using genome amplification approaches (Picogram input multimodal sequencing (PiMmS) method; Laumer (2023)), but these methods are yet to be widely employed by the biodiversity community.

### DNA SEQUENCE DATA

#### Reference sequences: specimen identification with DNA sequencing

DNA sequencing is a powerful tool for identifying specimens to species and genus levels, particularly when traditional morphological or biochemical methods are inadequate or inconclusive. Depending on the gene of interest, a minimum of one sequence (i.e. a single specimen) is recommended to define an OTU and sequencing multiple individuals from a population will provide an indication of diversity within a species. Obtaining sequences from multiple genes also increases confidence in the accuracy of the identification, as any sequencing errors (particularly relevant for Sanger sequencing since errors caused by nuclear copies of mitochondrial DNA (NUMTs are a possibility (Richly and Leister 2004)), or contamination can be detected and potentially corrected. Proper retention and storage of raw reads is important for verification and reproducibility of the data. Raw data, such as trace files, can be uploaded to BOLD alongside additional metadata (see below).

#### Primers

Primer mismatch is known to be a major issue in amplification of barcode fragments. Universal primers (LCOI490/HCO2198 (Folmer *et al*. 1994) and C1-J-1718 (Simon *et al*. 1994)) are effective for *COI* amplification for representatives of many animal phyla (Hebert *et al*. 2003). However, in agreement with past studies, we observed that these standard primers can amplify inconsistently across different groups in a variety of subterranean crustacean taxa (e.g. Camacho *et al*. 2011; Karanovic and Cooper 2012; Asmyhr and Cooper 2012). The universal primers for the nuclear *18S* (Whiting 2002) and mitochondrial *12S* rRNA (Wetzer 2001) genes are known to amplify well in many crustacean taxa (Asmyhr and Cooper 2012) and worked well here to produce alternative barcodes (Corse *et al*. 2010; Hardy *et al*. 2011). The standard *16S* primers (16sar/16Sbr Palumbi (1996)), whilst not successful for many taxa, were extremely useful for gastropods and copepods in this study. For these reasons, many studies develop *ad hoc* primers depending on the taxon group (e.g., Little *et al*. (2016)). In future, it will become increasingly feasible and affordable to sequence whole mitochondrial genomes employing next-generation sequencing (e.g., eDNA metagenomics), obtaining many barcoding genes at once or, alternatively, using the entire mitochondrial genome for species delineation or identification.

For a BRL to be useful for implementation into future eDNA metabarcoding studies, the barcodes of voucher specimens should match the most commonly used assay regions. Due to the historical technological impediment of the last 20 years where Sanger sequencing of barcode loci was the most inexpensive, subterranean fauna have most commonly been sequenced for loci including *COI*, *16S, 12S* and *18S,* in order of frequency. BRL DNA sequences should be homologous to metabarcoding assay regions (i.e., standard barcodes).

Currently, *COI* and *18S* are most commonly implemented for metazoan metabarcoding detection, and *16S* for microbial detection from eDNA. Commonly used primers for eDNA metabarcoding assays of freshwater invertebrates include *18S* (Pochon *et al*. 2013), *16S* (such as for crustaceans (Berry *et al*. 2017) and insects (Clarke *et al*. 2014)) and *COI* (Vamos *et al*. 2017). In future, genome skimming, including shotgun sequencing of mitochondrial genomes and Ultra Conserved Elements, may effectively be used as a universal approach for generating sequence data, especially for genuinely old and degraded specimens (Blaimer *et al*. 2016; McCormack *et al*. 2016) but also for contemporary specimens.

### METADATA

#### Linking specimen voucher information to DNA sequence and best available taxonomy

For an effective custom BRL database, the link between DNA sequences, morphologically identified specimens by a taxonomic practitioner (where possible), specimen metadata and the most up to date available taxonomy, which can be linked to sequence data are critical to its usability. A comprehensive guide to curation of a BRL for aquatic organisms (eukaryotic and prokaryotic) is described by Rimet *et al*. (2021) who recommended BOLD for its ability to link varied metadata to a sequence. For BOLD, minimum information per DNA sequence is: Sample ID (collection code), Field ID and/or Museum voucher ID (voucher number), Institution Storing, Phylum, and Country. However, it is strongly recommended that latitude and longitude (sample locality), morphological identification to lowest taxonomic level (Rimet *et al*. 2021), the name of the taxonomist, and the year of the identification also be included. BOLD allows submission of trace files for sequences and images (https://boldsystems.org/index.php/databases), which enhances accessibility and reusability. However, required information for GenBank is minimal and the infrastructure needed for hosting additional data, such as chromatograms and images, is not available, making it difficult to provide comprehensive metadata per specimen.

### DATABASE

#### Unified nomenclature and species delimitation

As mentioned previously, BOLD has developed the BIN system for *COI* sequences. The validity of using BINs as equivalent to species has been debated in the literature (Kekkonen *et al*. 2015; Meier *et al*. 2022). This is especially the case since use of *COI* only in conjunction with the OTU designation method of BOLD (i.e. the Refined Single Linkage (RESL) algorithm (Ratnasingham and Hebert 2013)), there is a tendency for over-splitting of lineages (Hlebec *et al*. 2023). However, we consider, as in other studies (e.g. Kekkonen *et al*. 2015; Blagoev *et al*. 2016), the utility of BINs as a built-in clustering method for generating species hypotheses is a valid approach, particularly for use by non-specialists. We observed that BINs, in conjunction with the ABGD species delimitation approach and gene fragment concordance to be consistent and in-line with the findings of other studies (Kekkonen *et al*. 2015). In future, the 29 lineages and 34 BINs we resolved here may be verified as undescribed species, with morphological examination of specimen vouchers.

GenBank, as the most widely used DNA sequence repository, can be used for querying metabarcoding reads and OTUs obtained in eDNA studies. Whilst this database is useful, especially given its centralization, the implementation and accessibility of metadata (e.g., taxonomic, geographic and photographic) associated with sequences means that there are added advantages to BOLD. Designed principally for biodiversity studies, the GUI of BOLD is uniquely configured to summarise geographic and taxonomic data. For example, phylogenetic trees can be directly estimated from DNA sequences, and the data can be stored in various projects and databases. Further, these aspects have the added benefit of being user-friendly for non-specialists, and provide an accessible implementation of the subterranean fauna BRL presented here. BOLD is also one of the most usable and collaborative databases available for depositing and analysing eDNA data. Whilst other databases, such as GenBank, have projects and databases (e.g. BioProjects, BioSamples, and the Sequence Read Archive), there are distinct advantages to using BOLD since, in addition to its ongoing funding, it is extremely attractive for the accessibility of custom BRLs for future research and, as such, we recommend its ongoing use for accessing our subterranean fauna database.

The Darwin Core Standard (DwC) offers a stable and flexible framework for compiling biodiversity data from varied and variable sources. Originally developed by the Biodiversity Information Standards community, Darwin Core is an evolving platform providing definitions and common terms that are shareable and searchable (Wieczorek *et al*. 2012). It plays a fundamental role in the sharing, use and reuse of open-access biodiversity data. We recommend that in curating data for custom BRL’s, the DwC standard terms be used. Importantly, BOLD, but not GenBank, implements the DwC standard (Ratnasingham and Hebert 2007).

#### Future taxonomic considerations

The usefulness of a subterranean fauna BRL rests in its ability to assign OTUs a unified nomenclature and, where possible, species names. Reference sequences must be directly linked to vouchered specimens of each species that are curated and maintained long-term within museum collections. For conservation management, species names are also critical (Costello *et al*., 2015), but as is the case for most invertebrate groups, especially subterranean fauna, formal taxonomy is not progressing rapidly enough compared to the recognition of the burgeoning number of molecular OTUs (Perina *et al*. 2018; Perina *et al*. 2019; Matthews *et al*. 2020). The number of new lineages we recovered in this study at Bungaroo Creek highlight the need for a unified nomenclature that is used consistently among stakeholders and to which taxonomy can be later applied, when available, to alleviate the well-documented taxonomic impediments (Riedel *et al*. 2013).

## Conclusions

A robust BRL is essential for the effective translation of anonymous DNA sequence data to meaningful taxonomic identification, including for the detection of fauna using metabarcoding. However, shareable, reusable and publicly available datasets for poorly known ecosystems are frequently unavailable for query. To address this lack of fundamental, but crucially important research, we have presented a custom BRL for an Australian subterranean fauna as a test case and, in doing so, outline best-practice standards for establishing a BRL to invigorate future subterranean faunal community assessment and monitoring. However, these recommendations also apply more broadly to all custom BRLs. Priorities to transform our consensus of species delimitation results into a more complete BRL in the future include morphological verification and taxonomic diagnoses of relevant species. A custom BRL requires DNA sequences at both broad (order, family) and narrow (genus to population level) taxonomic coverage to be useful in identifying unknown individuals and sequences in genetic and taxonomic studies (Hebert *et al*. 2013). Additional verification through more extensive sampling of subterranean fauna, barcoding of a much broader range of genes, and further phylogenetic analyses will also be essential. Given the uniquely diverse subterranean fauna present at Bungaroo Creek, and within the Pilbara region more broadly, a coordinated effort will be required to advance this data infrastructure in the future.

## Acknowledgements

This research was part funded by Australian Research Council (ARC) Industry Linkage Project Grant LP140100555 to A.D.A. S.J.B.C. and W.F.H. (linkage partners: South Australian Museum, Western Australian Museum, the WA Department of Biodiversity, Conservation and Attractions (DBCA) (formerly Department of Parks and Wildlife), Bennelongia Pty Ltd, Biota Environmental Sciences Pty Ltd), ARC Industry Linkage Project Grant LP190100555 to A.D.A. (linkage partners: Rio Tinto, BHP, Chevron, South Australian Museum, Western Australian Museum, WA DBCA, WA Department for Water and Environmental Regulation, Western Australian Biodiversity Science Institute (WABSI), Rio Tinto consulting contract to N.W. and M.B., and a research contract with WABSI funded by Rio Tinto and BHP. We thank S. Halse (Bennelongia) for collaboration and specimen collection as part of faunal surveys for ecological impact assessments, E. Matthews (formerly The University of Adelaide) for assistance collecting and sorting specimen samples in the Robe River Valley, Caitlin O’Neill, Dean Main and Michael Curran (Rio Tinto) for field support and valuable site information. Specimen processing and handling was undertaken at WAM by Ana Hara, Corey Whisson and Cathy Carr. Specimens were sampled under a Department of Parks and Wildlife Regulation 17 Licence to take fauna for scientific purposes (Number: 08-000700-1). This work was supported by resources provided by the Pawsey Supercomputing Centre with funding from the Australian Government and the Government of Western Australia.

## Notes

### Competing Interest Statement

The authors have declared no competing interest.

### Summary of Updates

Addition of two authors who were inadvertently not included in the original author list.

