## Supplementary Material 2 for "What are the best practices for curating eDNA custom barcode reference libraries? A case study using Australian subterranean fauna"

| Primers per gene | Annealing | Author |
| --- | --- | --- |
| *COI* |  |  |
| LCOI490 GGTCAACAAATCATAAAGATATTGG | 47-50ºC | (Folmer et al. 1994) |
| HCO2198 TGATTTTTTGGTCACCCTGAAGTTTA |  |  |
| *16S* |  |  |
| 16sar CGCCTGTTTATCAAAAAACAT |  | (Palumbi 1996) |
| 16Sbr ACGTGATCTGAGTTCAGACCGG |  |  |
| Nuclear *18S* |  |  |
| 1.2F TGCTTGTCTCAAAGATTAAGC | 49ºC | (Whiting 2002) |
| b5.0 TAACCGCAACAACTTTAAT (short) |  |  |
| b3.9 TGCTTTRAGCACTCTAA (long) |  | (Whiting et al. 1997) |
| Mitochondrial *12S* |  |  |
| 12SCRF GAGAGTGACGGGCGATATGT | 47ºC | (Wetzer 2001) |
| 12SCRR AAACCAGGATTAGATACCCTATTAT |  |  |

PCR-amplifications were conducted in 25 μL volumes comprising 1 x PCR buffer (1 mM dNTPs and 3 mM MgCl2), 1 unit of TAQ polymerase (MyTaq), approx. 1–5 ng of DNA template, and 0.2 μM of each primer. PCR cycling conditions as follows:

*COI* and *16S*: 5 min denaturing period at 95ºC, then 7 cycles of 95ºC for 30 s, 40ºC for 30 s, 72ºC for 60 s, followed by 35 cycles of 95ºC for 30 s, 50ºC for 30 s and 72ºC for 60 s with a final extension at 72ºC for 10 mins.

*18S* was amplified as three fragments, under the following conditions: 95ºC for 5 min, then 35 cycles of 95ºC for 30 s, 49ºC for 30 s, and 72ºC for 45 s, with a final extension of 72ºC for 10 mins.

*12S*: 95ºC for 5 min, then 40 cycles of 95ºC for 30 s, 47ºC for 30 s and 72ºC for 45 s, with a final extension at 72ºC for 10 mins. Minimal PCR optimisation was conducted.
