## Supplementary Material 3 for "What are the best practices for curating eDNA custom barcode reference libraries? A case study using Australian subterranean fauna"

| CO1 Stygofauna haplotypes (660bp) | | |  |  |  |  |  |  |  |  |  |
| --- | --- | --- | --- | --- | --- | --- | --- | --- | --- | --- | --- |
| Haplotype number | n | Sequences belonging to haplotype | BOLD BIN # | Group | | | | GENBANK ACCESSION | | | |
|  |  |  |  | CO1 | 18S | 12S | 16S | CO1 | 18S | 12S | 16S |
| 1 | 1 | WAMC72612 Pilbara sp.\|BUNWP0005 | BOLD:AEE8991 | 1 |  | 9 |  | OR524875 | | OR525169 | |
| 2 | 1 | WAMC71383 Amphipoda\|YALLWP01 | BOLD:AEE5579 | 2 |  | 9 |  | OR524819 | | OR525154 | |
| 3 | 2 | WAMC71128 Amphipoda\|BUNWB08 | BOLD:AEE7280 | 3 |  | 9 |  | OR524823 | | OR525158 | |
|  |  | WAMC71418 Amphipoda\|BUNWB08 | BOLD:AEE7280 | 3 | 1 | 9 |  | OR524813 | OR524935 | OR525150 | |
| 4 | 1 | WAMC72603 Nedsia sp.\|BUNWP0005 | BOLD:AEE7280 | 3 |  | 9 |  | OR524888 | | OR525196 | |
| 5 | 1 | WAMC72608 Nedsia sp.\|BUNWP0005 | BOLD:AEE7280 | 3 | 1 | 9 |  | OR524890 | OR525022 | OR525200 | |
| 6 | 4 | WAMC68601 Amphipoda\|JWO24 | BOLD:AEE9183 | 29 |  |  |  | OR524822 | |  |  |
|  |  | WAMC71104 Amphipoda\|JWO21 | BOLD:AEE9183 | 29 |  |  |  | OR524810 | |  |  |
|  |  | WAMC71130 Amphipoda\|BC225 | BOLD:AEE9183 | 29 | 1 | 9 |  | OR524806 | OR524930 | OR525144 | |
|  |  | WAMC71361 Amphipoda\|JWO21 | BOLD:AEE9183 | 29 |  |  |  | OR524811 | |  |  |
| 7 | 2 | WAMC71123 Amphipoda\|BC225 | BOLD:AEE9183 | 29 |  |  |  | OR524807 | |  |  |
|  |  | WAMC71124 Amphipoda\|BC225 | BOLD:AEE9183 | 29 |  |  |  | OR524817 | |  |  |
| 8 | 2 | WAMC71131 Amphipoda\|PZ11BUN008 | BOLD:AEE7196 | 29 |  | 9 |  | OR524809 | | OR525146 | |
|  |  | WAMC72610 Nedsia sp\|PZ11BUN008 | BOLD:AEE7196 | 29 | 1 | 9 |  | OR524889 | OR525019 | OR525197 | |
| 9 | 1 | WAMC68628 Amphipoda\|JWO24 | BOLD:AEE7196 | 29 |  |  |  | OR524818 | |  |  |
| 10 | 1 | WAMC71138 Amphipoda\|YALLWP01 | BOLD:AEE7196 | 29 |  | 9 |  | OR524821 | | OR525156 | |
| 11 | 1 | WAMT143820 Pauropodidae\|BC186 | BOLD:AEE8965 | 4 |  | 4 |  | OR524899 | | OR525226 | |
| 12 | 1 | WAMC71389 Ostracoda\|PZ11BUN008 | BOLD:AEE0612 | 5 | 11 | 6 |  | OR524893 | OR525031 | OR525211 | |
| 13 | 1 | WAMC71111 Ostracoda\|BC186 | BOLD:AEE1979 | 6 | 11 | 6 | 2 | OR524892 | OR525028 | OR525207 | OR524925 |
| 14 | 1 | WAMC71384 Ostracoda\|PZ11BUN008 | BOLD:AEE0613 | 7 | 11 | 6 |  | OR524891 | OR525024 | OR525203 | |
| 15 | 1 | WAMV9015 Tubificidae\|JWO23 | BOLD:AEE9005 | 8 | 14 | 2 | 9 | OR524887 | OR525016 | OR525193 | OR524924 |
| 16 | 1 | WAMV9014 Phreodrilidae\|BUNWO1105 | BOLD:AEE7132 | 9 | 14 | 1 | 7 | OR524900 | OR525041 | OR525227 | OR524927 |
| 17 | 1 | WAMV9023 Clitellata\|BUNWB09 | BOLD:AEE1055 | 10 | 14 | 3 | 6 | OR524827 | OR524944 | OR525167 | OR524911 |
| 18 | 1 | WAMV9012 Phreodrilidae\|BC225 | BOLD:AEE7131 | 11 | 14 | 3 | 4 | OR524902 | OR525043 | OR525229 | OR524929 |
| 19 | 1 | WAMV9013 Phreodrilidae\|BC667 | BOLD:AEE7562 | 12 | 14 | 3 | 5 | OR524901 | OR525042 | OR525228 | OR524928 |
| 20 | 1 | WAMC71404 Amphipoda\|PZ11BUN008 | BOLD:AEE8550 | 13 | 5 |  |  | OR524808 | OR524932 | |  |
| 21 | 1 | WAMC72616 Paramelitidae\|BC186 | BOLD:AEE9924 | 14 |  | 16 |  | OR524897 | | OR525223 | |
| 22 | 1 | WAMC72617 Paramelitidae\|BC186 | BOLD:AEE9924 | 14 |  | 16 |  | OR524898 | | OR525225 | |
| 23 | 1 | WAMC72619 Paramelitidae\|BC186 | BOLD:AEE9924 | 14 |  | 16 |  | OR524896 | | OR525221 | |
| 24 | 1 | WAMC71139 Amphipoda\|YALLWP01 | BOLD:AEE2761 | 15 | 2 |  |  | OR524814 | OR524936 | |  |
| 25 | 3 | WAMC71101 Amphipoda\|JWO24 | BOLD:AEE4462 | 16 |  | 15 |  | OR524820 | | OR525155 | |
|  |  | WAMC71363 Amphipoda\|JWO24 | BOLD:AEE4462 | 16 |  | 15 |  | OR524815 | | OR525152 | |
|  |  | WAMC71364 Amphipoda\|JWO24 | BOLD:AEE4462 | 16 |  |  |  | OR524812 | |  |  |
| 26 | 1 | WAMC68631 Amphipoda\|JWO24 | BOLD:AEE4462 | 16 |  |  |  | OR524816 | |  |  |
| 27 | 1 | WAMC72598 Bogidiella sp\|JWO24 | BOLD:AEE4462 | 16 |  | 15 |  | OR524826 | | OR525164 | |
| 28 | 1 | WAMC72596 Bogidiella sp\|JWO24 | BOLD:AEE4462 | 16 |  | 15 |  | OR524825 | | OR525161 | |
| 29 | 1 | WAMS71018 Gastropoda\|BUNWB09 | BOLD:AEE7100 | 17 | 13 | 8 | 3 | OR524886 | OR525008 | OR525184 | OR524919 |
| 30 | 2 | WAMC72710 Halosbae sp\|BC667 | BOLD:AEE4740 | 18 |  |  |  | OR524881 | |  |  |
|  |  | WAMC72712 Halosbae sp\|BC667 | BOLD:AEE4740 | 18 |  |  |  | OR524882 | |  |  |
| 31 | 1 | WAMC71100 Thermosbaecea\|JWO24 | BOLD:AEF0046 | 19 | 12 |  |  | OR524908 | OR525066 | |  |
| 32 | 1 | WAMC71372 Thermosbaecea\|PZ11BUN008 | BOLD:ADX1249 | 19 |  |  |  | OR524907 | |  |  |
| 33 | 1 | WAMC70445 Hexabathynella sp\|PZ10BUN004 | BOLD:AEE2039 | 20 |  | 10 |  | OR524885 | | OR525173 | |
| 34 | 1 | WAMC70449 nrBillibathynella\|BC292 | BOLD:AEE0507 | 21 | 17 | 14 |  |  |  |  |  |
| 35 | 1 | WAMC70447 Billibathynella\|BUNWP0006 | BOLD:AEE5164 | 22 | 17 | 14 |  | OR524824 | OR524941 | OR525160 | |
| 36 | 1 | WAMC70448 nrBillibathynella\|BUNWP0005 | BOLD:AEE8197 | 23 | 17 |  |  | OR524894 | OR525037 | |  |
| 37 | 2 | WAMC71419 Copepoda\|BC186 | BOLD:AEE5020 | 24 | 7 |  |  | OR524837 | OR524958 | |  |
|  |  | WAMC72627 Stygoridgewayia trispinosa\|BC186 | BOLD:AEE5020 | 24 | 7 |  |  | OR524905 | OR525050 | |  |
| 38 | 2 | WAMC71428 Copepoda\|BC186 | BOLD:AEE5020 | 24 | 7 |  |  | OR524847 | OR524973 | |  |
|  |  | WAMC72626 Stygoridgewayia trispinosa\|BC186 | BOLD:AEE5020 | 24 | 7 |  |  | OR524904 | OR525049 | |  |
| 39 | 1 | WAMC72625 Stygoridgewayia trispinosa\|BC186 | BOLD:AEE5020 | 24 | 7 |  |  | OR524903 | OR525048 | |  |
| 40 | 2 | WAMC71440 Copepoda\|JWO24 |  | 24 | 7 |  |  | OR524846 | OR524972 | |  |
|  |  | WAMC71459 Copepoda\|JWO24 |  | 24 | 7 |  |  | OR524854 | OR524979 | |  |
| 41 | 6 | WAMC71437 Copepoda\|JWO24 |  | 24 | 7 |  |  | OR524841 | OR524962 | |  |
|  |  | WAMC71448 Copepoda\|JWO24 |  | 24 | 7 |  |  | OR524843 | OR524965 | |  |
|  |  | WAMC71449 Copepoda\|JWO24 |  | 24 | 7 |  |  | OR524848 | OR524974 | |  |
|  |  | WAMC71452 Copepoda\|JWO24 |  | 24 |  |  |  | OR524852 | |  |  |
|  |  | WAMC71453 Copepoda\|JWO24 |  | 24 |  |  |  | OR524830 | |  |  |
|  |  | WAMC71460 Copepoda\|JWO24 |  | 24 | 7 |  |  | OR524853 | OR524978 | |  |
| 42 | 7 | WAMC71441 Copepoda\|JWO24 |  | 24 | 7 |  |  | OR524828 | OR524946 | |  |
|  |  | WAMC71444 Copepoda\|JWO24 |  | 24 | 7 |  |  | OR524856 | OR524981 | |  |
|  |  | WAMC71446 Copepoda\|JWO24 |  | 24 | 7 |  |  | OR524829 | OR524947 | |  |
|  |  | WAMC71450 Copepoda\|JWO24 |  | 24 | 7 |  |  | OR524855 | OR524980 | |  |
|  |  | WAMC71455 Copepoda\|JWO24 |  | 24 |  |  |  | OR524859 | |  |  |
|  |  | WAMC71457 Copepoda\|JWO24 |  | 24 | 7 |  |  | OR524845 | OR524971 | |  |
|  |  | WAMC71458 Copepoda\|JWO24 |  | 24 | 7 |  |  | OR524850 | OR524976 | |  |
| 43 | 4 | WAMC71442 Copepoda\|JWO24 |  | 24 | 7 |  |  | OR524857 | OR524982 | |  |
|  |  | WAMC71445 Copepoda\|JWO24 |  | 24 | 7 |  |  | OR524849 | OR524975 | |  |
|  |  | WAMC71447 Copepoda\|JWO24 |  | 24 | 7 |  |  | OR524858 | OR524983 | |  |
|  |  | WAMC71456 Copepoda\|JWO24 |  | 24 | 7 |  |  | OR524851 | OR524977 | |  |
| 44 | 3 | WAMC71425 Copepoda\|BC186 | BOLD:AEE5021 | 24 | 7 |  |  | OR524832 | OR524951 | |  |
|  |  | WAMC71426 Copepoda\|BC186 | BOLD:AEE5021 | 24 | 7 |  |  | OR524831 | OR524948 | |  |
|  |  | WAMC72624 Stygoridgewayia trispinosa\|BC186 | BOLD:AEE5021 | 24 | 7 |  |  | OR524906 | OR525051 | |  |
| 45 | 1 | WAMC71136 Copepoda\|PZ11BUN008 | BOLD:AEE2775 | 24 | 7 |  |  | OR524844 | OR524968 | |  |
| 46 | 1 | WAMC71398 Copepoda\|PZ11BUN008 | BOLD:AEE2775 | 24 | 7 |  |  | OR524842 | OR524963 | |  |
| 47 | 1 | WAMC71393 Copepoda\|PZ11BUN008 | BOLD:AEE2775 | 24 | 7 |  |  | OR524840 | OR524960 | |  |
| 48 | 1 | WAMC68490 Diacyclops sp\|BC360 | BOLD:AEE2774 | 25 | 9 |  |  | OR524868 | OR524991 | |  |
| 49 | 3 | WAMC71129 Copepoda\|BUNWB08 | BOLD:AEE2774 | 25 |  |  |  | OR524839 | |  |  |
|  |  | WAMC71413 Copepoda\|BC667 | BOLD:AEE2774 | 25 |  |  |  | OR524836 | |  |  |
|  |  | WAMC71415 Copepoda\|BUNWB08 | BOLD:AEE2774 | 25 | 9 |  |  | OR524838 | OR524959 | |  |
| 50 | 1 | WAMC71416 Copepoda\|BUNWO1108 | BOLD:AEE7353 | 26 |  |  |  | OR524835 | |  |  |
| 51 | 1 | WAMC68556 Diacyclops humphreysi\|BC225 | BOLD:AEE7353 | 26 | 9 |  |  | OR524865 | OR524988 | |  |
| 52 | 2 | WAMC72701 Diacyclops humphreysi humphreysi\|BUNWB09 | BOLD:AEE7353 | 26 | 9 |  |  | OR524866 | OR524989 | |  |
|  |  | WAMC72702 Diacyclops humphreysi humphreysi\|BUNWB09 | BOLD:AEE7353 | 26 | 9 |  |  | OR524867 | OR524990 | |  |
| 53 | 2 | WAMC68532 Harpactacoida\|BUNWB09 | BOLD:AEE7339 | 27 | 8 |  |  | OR524883 | OR525004 | |  |
|  |  | WAMC72639 Elaphoidella nrhumphreysi\|BUNWB09 | BOLD:AEE7339 | 27 | 8 |  |  | OR524871 | OR524995 | |  |
| 54 | 4 | WAMC68533 Harpactacoida\|BUNWB09 | BOLD:AEE7339 | 27 | 8 |  |  | OR524884 | OR525005 | |  |
|  |  | WAMC72638 Elaphoidella nrhumphreysi\|BUNWB09 | BOLD:AEE7339 | 27 | 8 |  |  | OR524869 | OR524993 | |  |
|  |  | WAMC72640 Elaphoidella nrhumphreysi\|BUNWB09 | BOLD:AEE7339 | 27 | 8 |  |  | OR524870 | OR524994 | |  |
|  |  | WAMC72641 Elaphoidella nrhumphreysi\|BUNWB09 | BOLD:AEE7339 | 27 | 8 |  |  | OR524872 | OR524996 | |  |
| 55 | 2 | WAMC72642 Elaphoidella humphreysi\|BUNWB13 | BOLD:AEE7339 | 27 |  |  |  | OR524874 | |  |  |
|  |  | WAMC72644 Elaphoidella humphreysi\|BUNWB13 | BOLD:AEE7339 | 27 | 8 |  |  | OR524873 | OR524997 | |  |
| 56 | 1 | WAMC71432 Copepoda\|BCP001 | BOLD:AEE7339 | 27 | 8 |  |  | OR524833 | OR524952 | |  |
| 57 | 1 | WAMC68495 Halicyclops (Rochacyclops) rochai\|PZ11BUN003 | BOLD:AEE3241 | 28 |  |  |  | OR524878 | |  |  |
| 58 | 7 | WAMC68526 Cyclopoida\|BUNWB09 | BOLD:AEE3240 | 28 |  |  |  | OR524860 | |  |  |
|  |  | WAMC68527 Cyclopoida\|BUNWB09 | BOLD:AEE3240 | 28 |  |  |  | OR524861 | |  |  |
|  |  | WAMC68529 Cyclopoida\|BUNWB09 | BOLD:AEE3240 | 28 |  |  |  | OR524862 | |  |  |
|  |  | WAMC71127 Copepoda\|BUNWO1108 | BOLD:AEE3240 | 28 |  |  |  | OR524834 | |  |  |
|  |  | WAMC72628 Halicyclops (Rochacyclops) rochai\|BUNWB09 | BOLD:AEE3240 | 28 | 10 |  |  | OR524876 | OR525000 | |  |
|  |  | WAMC72630 Halicyclops (Rochacyclops) rochai\|BUNWB09 | BOLD:AEE3240 | 28 | 10 |  |  | OR524877 | OR525001 | |  |
|  |  | WAMC72631 Halicyclops (Rochacyclops) rochai\|BUNWB09 | BOLD:AEE3240 | 28 | 10 |  |  | OR524880 | OR525003 | |  |
| 59 | 3 | WAMC68528 Cyclopoida\|BUNWB09 | BOLD:AEE3240 | 28 |  |  |  | OR524864 | |  |  |
|  |  | WAMC68530 Cyclopoida\|BUNWB09 | BOLD:AEE3240 | 28 |  |  |  | OR524863 | |  |  |
|  |  | WAMC72629 Halicyclops (Rochacyclops) rochai\|BUNWB09 | BOLD:AEE3240 | 28 | 10 |  |  | OR524879 | OR525002 | |  |

| 18S stygofauna haplotypes (1017bp) | | | Group | | | GB Accession No. | | |
| --- | --- | --- | --- | --- | --- | --- | --- | --- |
| Haplotype number | n | Sequences belonging to haplotype | 18S | 12S | 16S | 18S | 12S | 16S |
| 1 | 1 | WAMC71383 Amphipoda\|YALLWP01 | 1 |  |  | OR524939 | |  |
| 2 | 1 | WAMC72611 Pilbara sp\|BUNWP0005 | 1 |  |  | OR524999 | |  |
| 3 | 1 | WAMC71104 Amphipoda\|JWO21 | 1 |  |  | OR524933 | |  |
| 4 | 1 | WAMC72606 Nedsia sp\|BUNWP0005 | 1 |  |  | OR525017 | |  |
| 5 | 1 | WAMC72605 Nedsia sp\|BUNWP0005 | 1 |  |  | OR525021 | |  |
| 6 | 6 | WAMC71128 Amphipoda\|BUNWB08 | 1 |  |  | OR524940 | |  |
|  |  | WAMC71418 Amphipoda\|BUNWB08 |  |  |  |  |  |  |
|  |  | WAMC72604 Nedsia sp\|BUNWP0005 |  |  |  |  |  |  |
|  |  | WAMC72607 Nedsia sp\|BUNWP0005 |  |  |  |  |  |  |
|  |  | WAMC72608 Nedsia sp\|BUNWP0005 |  |  |  |  |  |  |
|  |  | WAMC72609 Nedsia sp\|BUNWP0005 |  |  |  |  |  |  |
| 7 | 4 | WAMC71123 Amphipoda\|BC225 | 1 |  |  | OR524931 | |  |
|  |  | WAMC71124 Amphipoda\|BC225 |  |  |  |  |  |  |
|  |  | WAMC71130 Amphipoda\|BC225 |  |  |  |  |  |  |
|  |  | WAMC71361 Amphipoda\|JWO21 |  |  |  |  |  |  |
| 8 | 1 | WAMC72610 Nedsia sp\|PZ11BUN008 | 1 |  |  | OR525019 | |  |
| 9 | 1 | WAMC71139 Amphipoda\|YALLWP01 | 2 |  |  | OR524936 | |  |
| 10 | 1 | WAMC72613 Pilbarus sp\|BUNWP0005 | 3 |  |  | OR525044 | |  |
| 11 | 1 | WAMC71362 Amphipoda\|JWO21 | 4 | 17 |  | OR524937 | OR525151 | |
| 12 | 1 | WAMC71404 Amphipoda\|PZ11BUN008 | 5 |  |  | OR524932 | |  |
| 13 | 1 | WAMV9019 Platyhelminthes\|JWO21 | 6 |  |  | OR525046 | |  |
| 14 | 2 | WAMV9026 Platyhelminthes\|JWO21 | 6 |  |  | OR525045 | |  |
|  |  | WAMV9028 Platyhelminthes\|JWO21 |  |  |  |  |  |  |
| 15 | 6 | WAMC68499 Copepoda\|BC186 | 7 |  |  | OR524949 | |  |
|  |  | WAMC68503 Copepoda\|BC186 |  |  |  |  |  |  |
|  |  | WAMC71398 Copepoda\|PZ11BUN008 |  |  |  |  |  |  |
|  |  | WAMC71450 Copepoda\|JWO24 |  |  |  |  |  |  |
|  |  | WAMC71458 Copepoda\|JWO24 |  |  |  |  |  |  |
|  |  | WAMC71460 Copepoda\|JWO24 |  |  |  |  |  |  |
| 16 | 1 | WAMC71426 Copepoda\|BC186 | 7 |  |  | OR524948 | |  |
| 17 | 3 | WAMC71136 Copepoda\|PZ11BUN008 | 7 |  |  | OR524968 | |  |
|  |  | WAMC71393 Copepoda\|PZ11BUN008 |  |  |  |  |  |  |
|  |  | WAMC71399 Copepoda\|PZ11BUN008 |  |  |  |  |  |  |
| 18 | 5 | WAMC71419 Copepoda\|BC186 | 7 |  |  | OR524958 | |  |
|  |  | WAMC71428 Copepoda\|BC186 |  |  |  |  |  |  |
|  |  | WAMC71430 Copepoda\|BC186 |  |  |  |  |  |  |
|  |  | WAMC71445 Copepoda\|JWO24 |  |  |  |  |  |  |
|  |  | WAMC71456 Copepoda\|JWO24 |  |  |  |  |  |  |
| 19 | 4 | WAMC71429 Copepoda\|BC186 | 7 |  |  | OR524966 | |  |
|  |  | WAMC71441 Copepoda\|JWO24 |  |  |  |  |  |  |
|  |  | WAMC71442 Copepoda\|JWO24 |  |  |  |  |  |  |
|  |  | WAMC71449 Copepoda\|JWO24 |  |  |  |  |  |  |
| 20 | 6 | WAMC68501 Copepoda\|BC186 |  |  |  |  |  |  |
|  |  | WAMC71437 Copepoda\|JWO24 |  |  |  |  |  |  |
|  |  | WAMC71440 Copepoda\|JWO24 |  |  |  |  |  |  |
|  |  | WAMC71447 Copepoda\|JWO24 |  |  |  |  |  |  |
|  |  | WAMC72624 Stygoridgewayia trispinosa\|BC186 |  |  |  |  |  |  |
|  |  | WAMC72625 Stygoridgewayia trispinosa\|BC186 |  |  |  |  |  |  |
| 21 | 2 | WAMC72626 Stygoridgewayia trispinosa\|BC186 |  |  |  |  |  |  |
|  |  | WAMC72627 Stygoridgewayia trispinosa\|BC186 |  |  |  |  |  |  |
| 22 | 1 | WAMC71459 Copepoda\|JWO24 | 7 |  |  | OR524979 | |  |
| 23 | 2 | WAMC68500 Copepoda\|BC186 | 7 |  |  | OR524957 | |  |
|  |  | WAMC71425 Copepoda\|BC186 |  |  |  |  |  |  |
| 24 | 1 | WAMC71446 Copepoda\|JWO24 | 7 |  |  | OR524947 | |  |
| 25 | 3 | WAMC71444 Copepoda\|JWO24 |  |  |  |  |  |  |
|  |  | WAMC71448 Copepoda\|JWO24 |  |  |  |  |  |  |
|  |  | WAMC71457 Copepoda\|JWO24 |  |  |  |  |  |  |
| 26 | 1 | WAMC68487 Cyclopoida\|YALLWP01 | 7 |  |  | OR524984 | |  |
| 27 | 1 | WAMC68548 Stygoridgewayia trispinosa\|JWO24 | 7 |  |  | OR525052 | |  |
| 28 | 1 | WAMC68505 Copepoda\|JWO24 | 7 |  |  | OR524961 | |  |
| 29 | 1 | WAMC71431 Copepoda\|BC186 | 7 |  |  | OR524970 | |  |
| 30 | 9 | WAMC68532 Harpactacoida\|BUNWB09 | 8 |  |  | OR525004 | |  |
|  |  | WAMC68533 Harpactacoida\|BUNWB09 |  |  |  |  |  |  |
|  |  | WAMC71432 Copepoda\|BCP001 |  |  |  |  |  |  |
|  |  | WAMC72638 Elaphoidella nr humphreysi\|BUNWB09 |  |  |  |  |  |  |
|  |  | WAMC72639 Elaphoidella nr humphreysi\|BUNWB09 |  |  |  |  |  |  |
|  |  | WAMC72640 Elaphoidella nr humphreysi\|BUNWB09 |  |  |  |  |  |  |
|  |  | WAMC72641 Elaphoidella nr humphreysi\|BUNWB09 |  |  |  |  |  |  |
|  |  | WAMC72643 Elaphoidella humphreysi\|BUNWB13 |  |  |  |  |  |  |
|  |  | WAMC72644 Elaphoidella humphreysi\|BUNWB13 |  |  |  |  |  |  |
| 31 | 1 | WAMC68572 Copepoda\|BUNWO1108 | 9 |  |  | OR524956 | |  |
| 32 | 1 | WAMC72703 Diacyclops humphreysi humphreysi\|BUNWB09 | 9 |  |  | OR524986 | |  |
| 33 | 5 | WAMC68488 Cyclopoida\|BUNWO1108 | 9 |  |  | OR524985 | |  |
|  |  | WAMC68492 Diacyclops sp\|BC360 |  |  |  |  |  |  |
|  |  | WAMC68556 Diacyclops humphreysi\|BC225 |  |  |  |  |  |  |
|  |  | WAMC72701 Diacyclops humphreysi humphreysi\|BUNWB09 |  |  |  |  |  |  |
|  |  | WAMC72702 Diacyclops humphreysi humphreysi\|BUNWB09 |  |  |  |  |  |  |
| 34 | 2 | WAMC68490 Diacyclops sp\|BC360 | 9 |  |  | OR524991 | |  |
|  |  | WAMC71415 Copepoda\|BUNWB08 |  |  |  |  |  |  |
| 35 | 1 | WAMC68524 Copepoda\|YALLWP01 | 9 |  |  | OR524969 | |  |
| 36 | 1 | WAMC68564 Diacyclops humphreysi\|BC292 | 9 |  |  | OR524987 | |  |
| 37 | 1 | WAMC71118 Copepoda\|BCP001 | 10 |  |  | OR524964 | |  |
| 38 | 5 | WAMC71427 Copepoda\|BC186 | 10 |  |  | OR524955 | |  |
|  |  | WAMC72628 Halicyclops (Rochacyclops) rochai\|BUNWB09 |  |  |  |  |  |  |
|  |  | WAMC72629 Halicyclops (Rochacyclops) rochai\|BUNWB09 |  |  |  |  |  |  |
|  |  | WAMC72630 Halicyclops (Rochacyclops) rochai\|BUNWB09 |  |  |  |  |  |  |
|  |  | WAMC72631 Halicyclops (Rochacyclops) rochai\|BUNWB09 |  |  |  |  |  |  |
| 39 | 1 | WAMC71385 Ostracoda\|PZ11BUN008 | 11 |  |  | OR525032 | |  |
| 40 | 7 | WAMC71109 Ostracoda\|JWO24 | 11 | 6 | 1 | OR525036 | OR525216 | OR524926 |
|  |  | WAMC71111 Ostracoda\|BC186 |  |  |  |  |  |  |
|  |  | WAMC71386 Ostracoda\|PZ11BUN008 |  |  |  |  |  |  |
|  |  | WAMC71389 Ostracoda\|PZ11BUN008 |  |  |  |  |  |  |
|  |  | WAMC71390 Ostracoda\|PZ11BUN008 |  |  |  |  |  |  |
|  |  | WAMC71410 Ostracoda\|JWO24 |  |  |  |  |  |  |
|  |  | WAMC71411 Ostracoda\|JWO24 |  |  |  |  |  |  |
| 41 | 5 | WAMC71384 Ostracoda\|PZ11BUN008 | 11 |  |  | OR525024 | |  |
|  |  | WAMC71387 Ostracoda\|PZ11BUN008 |  |  |  |  |  |  |
|  |  | WAMC71388 Ostracoda\|PZ11BUN008 |  |  |  |  |  |  |
|  |  | WAMC71391 Ostracoda\|PZ11BUN008 |  |  |  |  |  |  |
|  |  | WAMC71392 Ostracoda\|PZ11BUN008 |  |  |  |  |  |  |
| 42 | 1 | WAMC71368 Thermosbaecea\|PZ11BUN008 | 12 |  |  | OR525063 | |  |
| 43 | 2 | WAMC71405 Thermosbaecea\|BC186 | 12 |  |  | OR525060 | |  |
|  |  | WAMC71408 Thermosbaecea\|BC186 |  |  |  |  |  |  |
| 44 | 1 | WAMC71407 Thermosbaecea\|BC186 | 12 |  |  | OR525053 | |  |
| 45 | 1 | WAMC71100 Thermosbaecea\|JWO24 | 12 |  |  | OR525066 | |  |
| 46 | 9 | WAMC71110 Thermosbaecea\|BC186 | 12 |  |  | OR525059 | |  |
|  |  | WAMC71134 Thermosbaecea\|PZ11BUN008 |  |  |  |  |  |  |
|  |  | WAMC71365 Thermosbaecea\|JWO24 |  |  |  |  |  |  |
|  |  | WAMC71366 Thermosbaecea\|PZ11BUN008 |  |  |  |  |  |  |
|  |  | WAMC71367 Thermosbaecea\|PZ11BUN008 |  |  |  |  |  |  |
|  |  | WAMC71370 Thermosbaecea\|PZ11BUN008 |  |  |  |  |  |  |
|  |  | WAMC71371 Thermosbaecea\|PZ11BUN008 |  |  |  |  |  |  |
|  |  | WAMC71380 Thermosbaecea\|PZ11BUN008 |  |  |  |  |  |  |
|  |  | WAMC71409 Thermosbaecea\|BC186 |  |  |  |  |  |  |
| 47 | 5 | WAMS71012 Gastropoda\|BUNWB13 | 13 |  |  | OR525010 | |  |
|  |  | WAMS71014 Gastropoda\|BUNWB08 |  |  |  |  |  |  |
|  |  | WAMS71018 Gastropoda\|BUNWB09 |  |  |  |  |  |  |
|  |  | WAMS71019 Gastropoda\|BUNWB09 |  |  |  |  |  |  |
|  |  | WAMS71024 Gastropoda\|BUNWB13 |  |  |  |  |  |  |
| 48 | 1 | WAMV9015 Tubificidae\|JWO23 | 14 |  |  | OR525016 | |  |
| 49 | 2 | WAMV9020 Clitellata\|JWO21 | 14 |  |  | OR524945 | |  |
|  |  | WAMV9025 Clitellata\|JWO21 |  |  |  |  |  |  |
| 50 | 1 | WAMV9022 Clitellata\|PZ11BUN008 | 14 |  |  | OR524943 | |  |
| 51 | 1 | WAMV9014 Phreodrilidae\|BUNWO1105 | 14 |  |  | OR525041 | |  |
| 52 | 1 | WAMV9023 Clitellata\|BUNWB09 | 14 |  |  | OR524944 | |  |
| 53 | 1 | WAMV9012 Phreodrilidae\|BC225 | 14 |  |  | OR525043 | |  |
| 54 | 1 | WAMV9013 Phreodrilidae\|BC667 | 14 |  |  | OR525042 | |  |
| 55 | 4 | WAMT144601 Lophoturus madecassus\|BC186 | 15 | 12 |  | OR525015 | OR525192 | |
|  |  | WAMT144602 Lophoturus madecassus\|BC186 |  |  |  |  |  |  |
|  |  | WAMT144603 Lophoturus madecassus\|BC186 |  |  |  |  |  |  |
|  |  | WAMT144604 Lophoturus madecassus\|BC186 |  |  |  |  |  |  |
| 56 | 1 | WAMC70440 Kagala tonde\|BUNWP0005 | 16 | 5 |  | OR525006 | OR525176 | |
| 57 | 2 | WAMC70447 Billibathynella sp\|BUNWP0006 | 17 |  |  | OR524941 | |  |
|  |  | WAMC70448 nrBillibathynella sp\|BUNWP0005 |  |  |  |  |  |  |
| 58 | 3 | WAMC70449 nrBillibathynella sp\|BC292 | 17 |  |  |  |  |  |
|  |  | WAMC70450 nrBillibathynella sp\|BUNWB13 |  | 13 |  |  | OR525218 | |
|  |  | WAMC72623 nrBillibathynella sp\|BC292 |  |  |  |  |  |  |

| 12S Stygofauna (425 bp) | | | Group | | GBAccNo | |
| --- | --- | --- | --- | --- | --- | --- |
| Hap. number | n | Sequences belonging to haplotype | 12S | 16S | 12S | 16S |
| 1 | 1 | WAMV9014 Phreodrilidae\|BUNWO1105 | 1 |  | OR525227 |  |
| 2 | 1 | WAMV9022 Clitellata\|PZ11BUN008 | 2 | 8 | OR525166 | OR524910 |
| 3 | 1 | WAMV9015 Tubificidae\|JWO23 | 2 |  | OR525193 |  |
| 4 | 2 | WAMV9020 Clitellata\|JWO21 | 2 |  | OR525168 |  |
|  |  | WAMV9025 Clitellata\|JWO21 |  |  |  |  |
| 5 | 1 | WAMV9023 Clitellata\|BUNWB09 | 3 |  | OR525167 |  |
| 6 | 1 | WAMV9012 Phreodrilidae\|BC225 | 3 |  | OR525229 |  |
| 7 | 1 | WAMV9013 Phreodrilidae\|BC667 | 3 |  | OR525228 |  |
| 8 | 1 | WAMT143820 Pauropodidae\|BC186 | 4 |  | OR525226 |  |
| 9 | 1 | WAMC70439 Kagala tonde\|JWO23 | 5 |  | OR525177 |  |
| 10 | 1 | WAMC70440 Kagala tonde\|BUNWP0005 | 5 |  | OR525176 |  |
| 11 | 2 | WAMC71109 Ostracoda\|JWO24 | 6 |  | OR525216 |  |
|  |  | WAMC71410 Ostracoda\|JWO24 |  |  |  |  |
| 12 | 1 | WAMC71411 Ostracoda\|JWO24 | 6 |  | OR525210 |  |
| 13 | 1 | WAMC71389 Ostracoda\|PZ11BUN008 | 6 |  | OR525211 |  |
| 14 | 3 | WAMC71384 Ostracoda\|PZ11BUN008 | 6 |  | OR525203 |  |
|  |  | WAMC71387 Ostracoda\|PZ11BUN008 |  |  |  |  |
|  |  | WAMC71388 Ostracoda\|PZ11BUN008 |  |  |  |  |
| 15 | 1 | WAMC71385 Ostracoda\|PZ11BUN008 | 6 |  | OR525212 |  |
| 16 | 2 | WAMC71391 Ostracoda\|PZ11BUN008 | 6 |  | OR525208 |  |
|  |  | WAMC71392 Ostracoda\|PZ11BUN008 |  |  |  |  |
| 17 | 2 | WAMC71137 Ostracoda\|PZ11BUN008 |  |  |  |  |
|  |  | WAMC71386 Ostracoda\|PZ11BUN008 |  |  |  |  |
| 18 | 1 | WAMC71390 Ostracoda\|PZ11BUN008 | 6 |  | OR525204 |  |
| 19 | 2 | WAMC70516 Humphreyscando sp\|JWO24 | 6 |  | OR525174 |  |
|  |  | WAMC72664 Humphreyscando sp\|BC405 |  |  |  |  |
| 20 | 1 | WAMC71111 Ostracoda\|BC186 | 6 |  | OR525207 |  |
| 21 | 1 | WAMC71406 Thermosbaecea\|BC186 | 7 |  | OR525231 |  |
| 22 | 9 | WAMS71011 Gastropoda\|BUNWB13 | 8 |  | OR525187 |  |
|  |  | WAMS71012 Gastropoda\|BUNWB13 |  |  |  |  |
|  |  | WAMS71014 Gastropoda\|BUNWB08 |  |  |  |  |
|  |  | WAMS71015 Gastropoda\|BUNWB08 |  |  |  |  |
|  |  | WAMS71019 Gastropoda\|BUNWB09 |  |  |  |  |
|  |  | WAMS71020 Gastropoda\|BUNWB09 |  |  |  |  |
|  |  | WAMS71022 Gastropoda\|BUNWB13 |  |  |  |  |
|  |  | WAMS71023 Gastropoda\|BUNWB13 |  |  |  |  |
|  |  | WAMS71024 Gastropoda\|BUNWB13 |  |  |  |  |
| 23 | 2 | WAMS71017 Gastropoda\|BUNWB09 | 8 |  | OR525180 |  |
|  |  | WAMS71018 Gastropoda\|BUNWB09 |  |  |  |  |
| 24 | 1 | WAMC71383 Amphipoda\|YALLWP01 | 9 |  | OR525154 |  |
| 25 | 1 | WAMC72611 Pilbara sp\|BUNWP0005 | 9 |  | OR525170 |  |
| 26 | 1 | WAMC72612 Pilbara sp\|BUNWP0005 | 9 |  | OR525169 |  |
| 27 | 11 | WAMC71128 Amphipoda\|BUNWB08 | 9 |  | OR525158 |  |
|  |  | WAMC71418 Amphipoda\|BUNWB08 |  |  |  |  |
|  |  | WAMC72601 Nedsia sp\|BUNWP0005 |  |  |  |  |
|  |  | WAMC72603 Nedsia sp\|BUNWP0005 |  |  |  |  |
|  |  | WAMC72604 Nedsia sp\|BUNWP0005 |  |  |  |  |
|  |  | WAMC72605 Nedsia sp\|BUNWP0005 |  |  |  |  |
|  |  | WAMC72606 Nedsia sp\|BUNWP0005 |  |  |  |  |
|  |  | WAMC72607 Nedsia sp\|BUNWP0005 |  |  |  |  |
|  |  | WAMC72608 Nedsia sp\|BUNWP0005 |  |  |  |  |
|  |  | WAMC72609 Nedsia sp\|BUNWP0005 |  |  |  |  |
| 28 | 3 | WAMC71131 Amphipoda\|PZ11BUN008 | 9 |  | OR525146 |  |
|  |  | WAMC71382 Amphipoda\|YALLWP01 |  |  |  |  |
|  |  | WAMC72610 Nedsia sp\|PZ11BUN008 |  |  |  |  |
| 29 | 5 | WAMC71104 Amphipoda\|JWO21 | 9 |  | OR525147 |  |
|  |  | WAMC71123 Amphipoda\|BC225 |  |  |  |  |
|  |  | WAMC71124 Amphipoda\|BC225 |  |  |  |  |
|  |  | WAMC71130 Amphipoda\|BC225 |  |  |  |  |
|  |  | WAMC71361 Amphipoda\|JWO21 |  |  |  |  |
| 30 | 1 | WAMC71138 Amphipoda\|YALLWP01 | 9 |  | OR525156 |  |
| 31 | 1 | WAMC70446 Hexabathynella sp\|BC401 | 10 |  | OR525172 |  |
| 32 | 1 | WAMC70445 Hexabathynella sp\|PZ10BUN004 | 10 |  | OR525173 |  |
| 33 | 1 | WAMC72621 Hexabathynella sp\|PZ10BUN004 | 10 |  | OR525171 |  |
| 34 | 1 | WAMC71378 Thermosbaecea\|PZ11BUN008 | 11 |  | OR525230 |  |
| 35 | 4 | WAMT144601 Lophoturus madecassus\|BC186 | 12 |  | OR525192 |  |
|  |  | WAMT144602 Lophoturus madecassus\|BC186 |  |  |  |  |
|  |  | WAMT144603 Lophoturus madecassus\|BC186 |  |  |  |  |
|  |  | WAMT144604 Lophoturus madecassus\|BC186 |  |  |  |  |
| 36 | 1 | WAMC70450 nrBillibathynella sp\|BUNWB13 | 13 |  | OR525218 |  |
| 37 | 1 | WAMC70447 Billibathynella sp\|BUNWP0006 | 14 |  | OR525160 |  |
| 38 | 1 | WAMC70449 nrBillibathynella sp\|BC292 | 14 |  |  |  |
| 39 | 4 | WAMC71101 Amphipoda\|JWO24 | 15 |  | OR525155 |  |
|  |  | WAMC71363 Amphipoda\|JWO24 |  |  |  |  |
|  |  | WAMC71364 Amphipoda\|JWO24 |  |  |  |  |
|  |  | WAMC72597 Bogidiella sp\|JWO24 |  |  |  |  |
| 40 | 2 | WAMC72596 Bogidiella sp\|JWO24 | 15 |  | OR525161 |  |
|  |  | WAMC72598 Bogidiella sp\|JWO24 |  |  |  |  |
| 41 | 1 | WAMC72600 Bogidiella sp\|JWO24 | 15 |  | OR525163 |  |
| 42 | 1 | WAMC72616 Paramelitidae\|BC186 | 16 |  | OR525223 |  |
| 43 | 1 | WAMC72619 Paramelitidae\|BC186 | 16 |  | OR525221 |  |
| 44 | 1 | WAMC72617 Paramelitidae\|BC186 | 16 |  | OR525225 |  |
| 45 | 2 | WAMC70433 Paramelitidae\|BC225 | 17 |  | OR525219 |  |
|  |  | WAMC72614 Paramelitidae\|BC186 |  |  |  |  |
| 46 | 1 | WAMC71362 Amphipoda\|JWO21 | 17 |  | OR525151 |  |
| 47 | 1 | WAMC71112 Amphipoda\|BC186 | 17 |  | OR525157 |  |
| 48 | 2 | WAMC72615 Paramelitidae\|BC186 | 17 |  | OR525224 |  |
|  |  | WAMC72618 Paramelitidae\|BC186 |  |  |  |  |

| 16S stygofauna (511 bp) | | | Group | GBAccNo |
| --- | --- | --- | --- | --- |
| Hap. Number | n | Sequences belonging to haplotype | 16S | 16s |
| 1 | 1 | WAMC71109 Ostracoda\|JWO24 | 1 | OR524926 |
| 2 | 1 | WAMC71111 Ostracoda\|BC186 | 2 | OR524925 |
| 3 | 9 | WAMS71011 Gastropoda\|BUNWB13 | 3 | OR524922 |
|  |  | WAMS71012 Gastropoda\|BUNWB13 |  |  |
|  |  | WAMS71014 Gastropoda\|BUNWB08 |  |  |
|  |  | WAMS71015 Gastropoda\|BUNWB08 |  |  |
|  |  | WAMS71017 Gastropoda\|BUNWB09 |  |  |
|  |  | WAMS71018 Gastropoda\|BUNWB09 |  |  |
|  |  | WAMS71022 Gastropoda\|BUNWB13 |  |  |
|  |  | WAMS71023 Gastropoda\|BUNWB13 |  |  |
|  |  | WAMS71024 Gastropoda\|BUNWB13 |  |  |
| 4 | 2 | WAMS71019 Gastropoda\|BUNWB09 | 3 | OR524914 |
|  |  | WAMS71020 Gastropoda\|BUNWB09 |  |  |
| 5 | 1 | WAMV9012 Phreodrilidae\|BC225 | 4 | OR524929 |
| 6 | 1 | WAMV9013 Phreodrilidae\|BC667 | 5 | OR524928 |
| 7 | 1 | WAMV9023 Clitellata\|BUNWB09 | 6 | OR524911 |
| 8 | 1 | WAMV9014 Phreodrilidae\|BUNWO1105 | 7 | OR524927 |
| 9 | 1 | WAMV9022 Clitellata\|PZ11BUN008 | 8 | OR524910 |
| 10 | 1 | WAMV9015 Phreodrilidae\|JWO23 | 9 | OR524924 |
| 11 | 2 | WAMV9020 Clitellata\|JWO21 | 9 | OR524912 |
|  |  | WAMV9025 Clitellata\|JWO21 |  |  |
