## Supplementary Material 4 for "What are the best practices for curating eDNA custom barcode reference libraries? A case study using Australian subterranean fauna"

**Supplementary Material 4:** Phylogenetic trees of taxa from Bungaroo creek for a) *18S* rRNA, b) *12S* rRNA, c) *16S* rRNA – with species delimitation results for represented by taxonomic group in colour: Amphipoda (blue), Copepoda (purple), Bathynellacea (green), Thermosbaenacea (pink), Ostracoda (red), Annelida (orange), Gastropoda (blue). Relationships between major taxonomic groups are not representative of phylogenetic relationships.

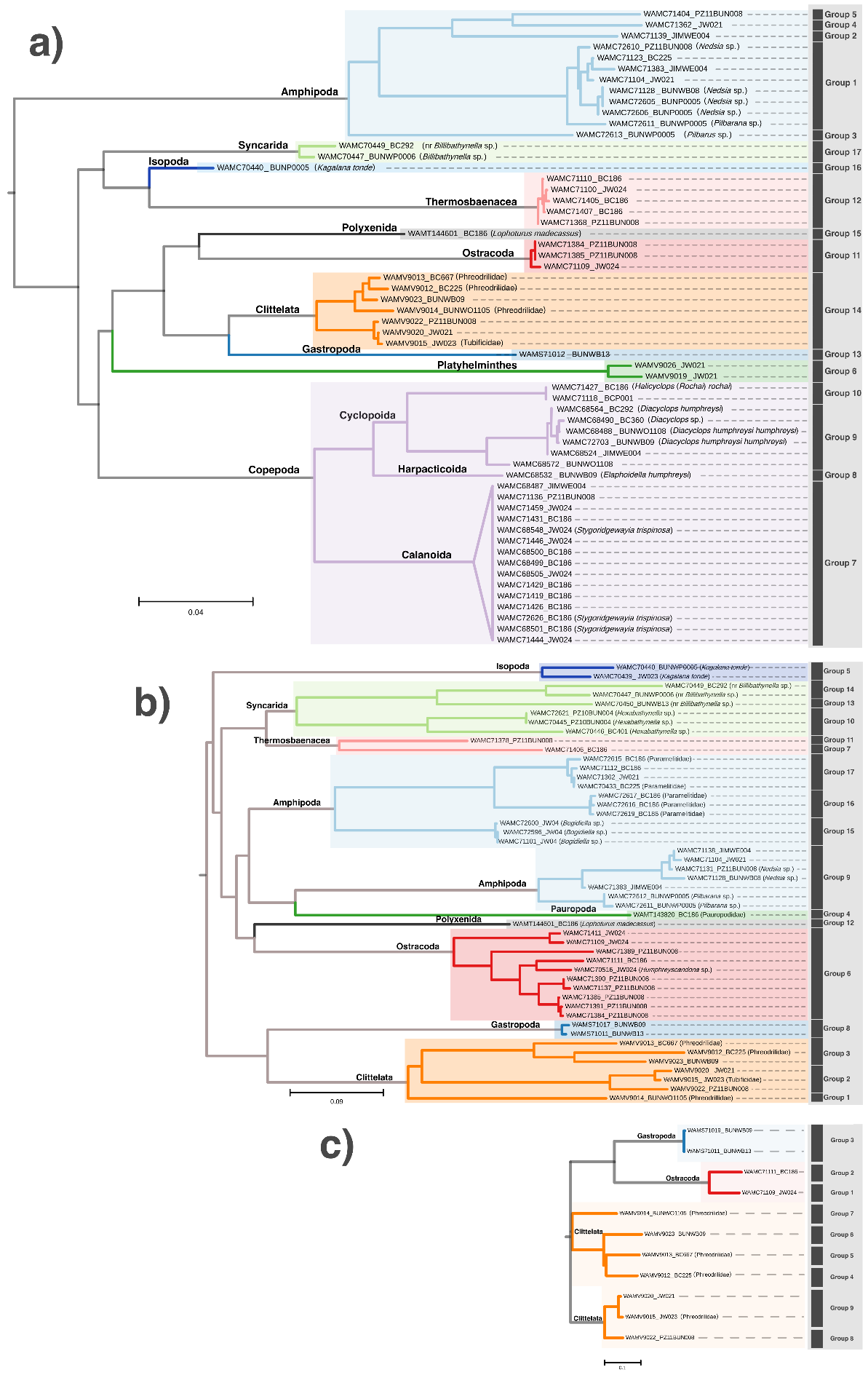
